# Cholesterol esterification blockade boosts immunotherapy and reduces cancer relapse

**DOI:** 10.1101/2025.10.15.682531

**Authors:** Yange Gu, Lulu Zhang, Jianhua Li, Ensi Ma, Weiwei Hu, Xiaocheng Du, Feifei Zhang, Tiange Yang, Linyue Bian, Yitian Jin, Yun Liu, Yinghua Wang, Zihan Ning, Chune Yu, Dun Li, Jing Zhao, Chenqi Xu, Zhengxin Wang, Guangchuan Wang

## Abstract

Recurrence after liver cancer resection and transplantation remains a critical clinical challenge, driven by immune evasion. However, effective immune-sensitizing targets for preventing or managing relapse remain limited. In this study, we integrated differential proteomic profiling of human liver cancer samples with or without subsequent recurrence and T cell killing assays to uncover a pivotal role for the cholesterol esterification enzyme SOAT1 in immune evasion and cancer recurrence. Genetic knockout or pharmacological inhibition of SOAT1 markedly sensitized both mouse and human cancer cells to immune surveillance and T cell-mediated killing. Mechanistically, blocking cholesterol esterification disrupted excessive cholesterol biosynthesis, impairing cancer cells’ antioxidant capacity and metabolic resilience under immune attack. Importantly, SOAT1 inhibition effectively suppressed tumor immune escape and improved the efficacy of anti-PD1 and CAR-T cell therapies, even in immunocompromised conditions. These findings highlight cholesterol esterification as a key driver of cancer cell redox metabolism resilience under immunosurveillance and position SOAT1 as a promising target to prevent cancer relapse and enhance immunotherapy outcomes.

## Introduction

Liver transplantation is one of the most effective treatments for hepatocellular carcinoma (HCC) patients who are ineligible for resection or suffer from decompensated cirrhosis^1^. However, post-transplant tumor recurrence remains a major challenge, with rates as high as 40%, severely impairing patient prognosis^2,3^. Furthermore, recurrence rates after hepatic resection can reach up to 70% within five years^1^. Despite the high prevalence of recurrence, there are currently no specific targets to effectively prevent HCC recurrence or manage relapsed tumors^1,4^.

Immunosurveillance is crucial for preventing cancer recurrence and metastases. However, cancer cells often hijack inherent or altered metabolic pathways to evade immune destruction and resist therapies, facilitating recurrence^3–7^. This is particularly relevant in HCC, where metabolic disorders—especially those associated with obesity and hypercholesterolemia—are key drivers of tumor progression and recurrence^8–11^. Non-alcoholic fatty liver disease (NAFLD) and obesity-associated metabolic dysfunctions have emerged as critical contributors to both liver cancer development and recurrence^14–16^. While immunotherapies, such as immune checkpoint inhibitors, have revolutionized cancer treatment with remarkable success in many cancer types^14–16^, the response rates in HCC remain disappointingly low^17^. This highlights the urgent need to identify liver cancer-specific drivers and mechanisms of immune evasion and recurrence, to enable more effective therapeutic approaches to prevent recurrence and manage relapsed tumors.

In this study, we utilized proteomics analysis of liver cancer samples with and without recurrence, followed by T cell killing assays, to identify sterol O-acyltransferase 1 (SOAT1) as a key driver of immune evasion and cancer recurrence in HCC. Targeting SOAT1 through genetic knockout or chemical inhibitors significantly enhanced the sensitivity of liver cancer cells to T cell-mediated killing and anti-PD1 immunotherapy, while reducing tumor growth and recurrence under both normal and compromised immune conditions. Mechanistically, SOAT1 knockout led to cholesterol accumulation in the endoplasmic reticulum (ER), disrupting SREBP1/2-mediated cholesterol and fatty acids biosynthesis, thereby reducing the antioxidant capacity and metabolic resilience of cancer cells during T cell-mediated cytotoxicity. These findings highlight SOAT1 as a promising target for enhancing immunotherapy and preventing liver cancer recurrence, even in immunocompromised settings.

## Results

### Integrated proteomics and T cell killing assays identify SOAT1 as a key driver of immune evasion and recurrence in liver cancer

To identify key drivers of immune evasion and recurrence in liver cancer, we collected liver cancer samples from 28 liver transplant patients with HCC who received similar treatments but exhibited distinct recurrence patterns (**Figure 1A**). Based on follow-up data over 5 years, the patients were divided into two groups: a recurrence group of 15 patients with a median progression-free survival (PFS) of 346.8 days and a non-recurrence group of 13 patients with a PFS exceeding 1200 days, both groups having comparable baseline characteristics (**Figures 1B, 1C, S1A, and Table S1**). We then performed proteomics analysis on these primary HCC samples using liquid chromatography mass spectrometry (LC-MS), and identified 86 significantly differentially expressed proteins (DEPs) which may contribute to liver cancer recurrence (**Figures 1D and S1B**). Gene ontology (GO) analysis indicated upregulation of genes related to cholesterol metabolism, nucleotide excision repair, steroid hormone biosynthesis, and DNA replication in recurrent cancers, suggesting their role in liver cancer recurrence (**Figure 1E**).

**Figure 1.**
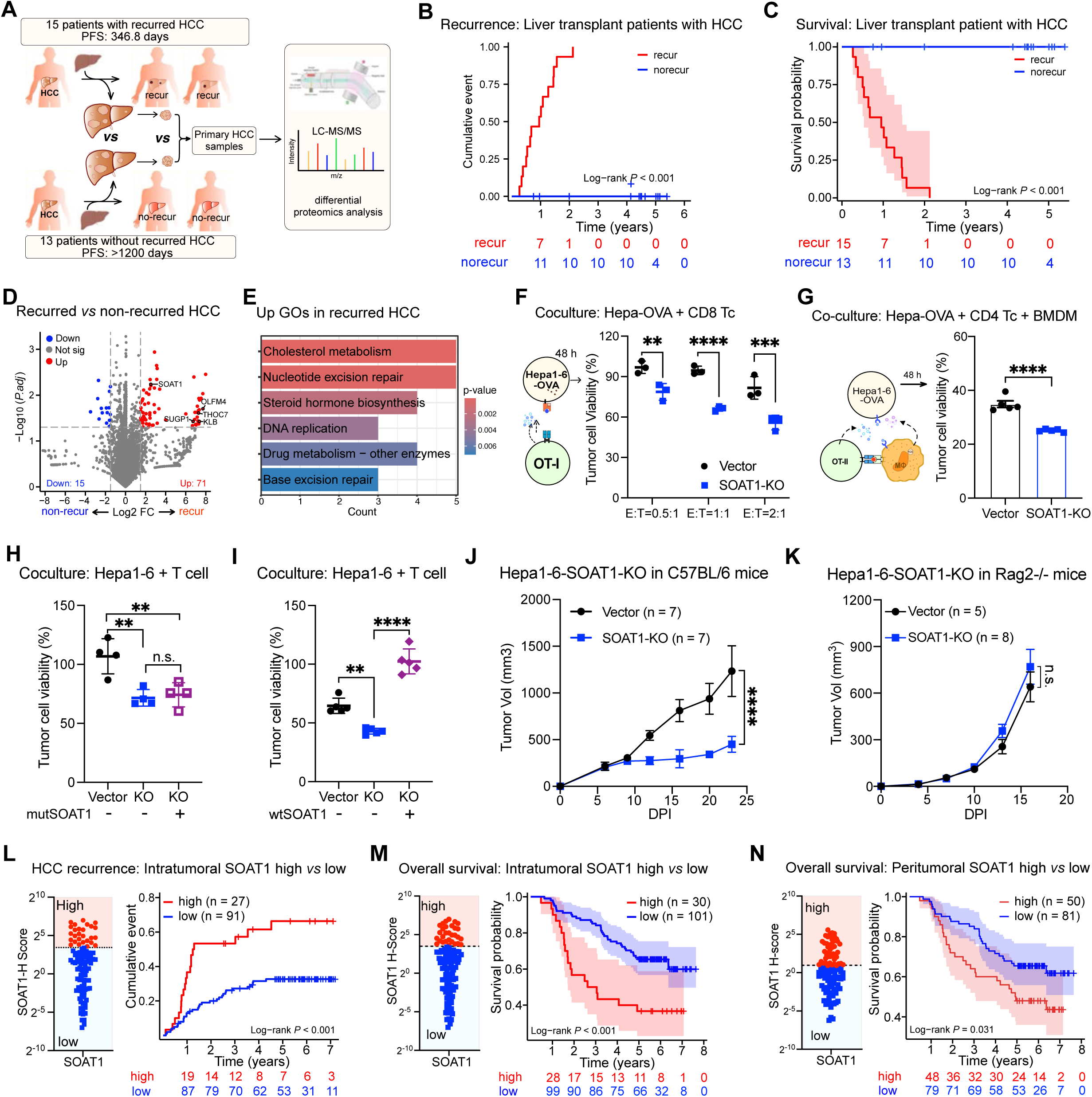
Proteomics analysis of recurrent versus nonrecurrent human HCC samples, combined with immune cell killing assays, identifies SOAT1 as a key driver of HCC immune evasion and recurrence. (**A**) Schematics illustrating the experimental design of collecting primary human HCC samples from liver transplant patients with HCC with later recurrence versus those without recurrence for proteomics analysis using liquid chromatography mass spectrometry (LC-MS). (**B, C**) Cumulative liver cancer recurrence curves (**B**) and Kaplan Meier survival curves (**C**) of 28 liver transplant patients with HCC. The recurrence group of 15 people had a medium PFS of 346.8 days, while the non-recurrence group had a medium PFS exceeding 1200 days.(**D**) Volcano plot of proteomics data depicting the differentially expressed proteins (DEPs, log2 FC ≥ 1.5 and P < 0.05) in the primary HCC tumors with later recurrence *versus* those without later recurrence. (**E**) GO analysis on the significantly upregulated DEPs in the primary HCCs with later recurrence versus those without later recurrence. (**F**) Viability of SOAT1-KO and vector control Hepa1-6-OVA cells after coculturing with OT-I CD8+T cells at E:T ratios of 0.5:1, 1:1, or 2:1 for 48 h (n = 3). (**G**) Viability of SOAT1-KO and vector control Hepa1-6-OVA cells after coculture with bone marrow-derived macrophages (BMDMs) at ratios of BMDMs: Hepa-OVA-Cas9 of 1:1, or OT-II CD4+T cells and BMDMs at ratios of CD4: BMDM: Hepa-OVA-Cas9 of 2:1:1 for 48 h (n = 3). (**H, I**) Viability of Hepa-OVA-Cas9 cells with vector, SOAT1-KO, and wildtype SOAT1-rescued (**H**), or catalytic-inactive mutant SOAT1-rescued (**I**) after coculture with OT-I CD8+ T cells at an E:T ratio of 2:1 for 48 h (n = 4). (**J, K**) Growth curves of subcutaneously inoculated SOAT1-KO and vector control Hepa1-6 tumors in immunodeficient Rag2^−/-^ mice (n = 5 for vector, n = 8 for SOAT1-KO) (**J**) and immunocompetent C57BL/6 mice (n = 7) (**K**). (**L, M**) Prognostic significance of intratumoral SOAT1 expression in HCC recurrence (**L**) and overall survival (**M**) in liver transplant patients was evaluated by stratifying patients into SOAT1-high (n = 30) and SOAT1-low (n = 101) groups. The SOAT1-high cohort exhibited a significantly higher recurrence rate (**L**) and poorer overall survival (**M**) compared to the SOAT1-low cohort. (**N**) Prognostic impact of peritumoral SOAT1 expression in liver transplant patients with HCC was assessed by stratifying patients into high (n = 50) and low peritumoral SOAT1 (n = 81). High peritumoral SOAT1 was associated with significantly worse overall survival. Data are shown as means ± SEMs; Statistical analysis was performed using a Student’s t test or two-way ANOVA. Asterisks: * P < 0.05, ** P < 0.01, *** P < 0.001, ****P <0.0001. See also: Supplementary Figures S1-S2 and Tables S1.

To investigate the contributions of cholesterol metabolism-related DEPs in liver cancer immune evasion and recurrence, we knocked out *Thoc7*, *Soat1*, *Sugp1*, *Olfm4*, *Klb* in cholesterol metabolism, as well as *Nyfa* and *Pole* individually in Hepa1-6 mouse liver cancer cells stably expressing Cas9 and OVA (Hepa-OVA-Cas9). Firstly, we assessed their roles in tumor cell proliferation and found that individual knockout of these genes had minimal impact on Hepa1-6 proliferation (**Figures S1C and S1D**). Next, to evaluate their roles in immune evasion, we cocultured them with antigen-specific OT-I CD8+ T cells. Knockout of *Pole*, *Sugp1*, and *Soat1* significantly increased the vulnerability of Hepa1-6 cells to T cell-mediated killing (**Figure S1E**), with *Soat1* knockout (SOAT1-KO) showing the most pronounced sensitivity (**Figure 1F**). Additionally, coculture of SOAT1-KO and vector Hepa-OVA-Cas9 cells with bone marrow-derived macrophages (BMDMs) and antigen specific OT-II CD4^+^ T cells demonstrated that SOAT1-KO cells were more susceptible to inflammatory killing^21^ (**Figure 1G**). Moreover, in another C57BL/6 syngeneic HCC cell line, HCC51 (CAG-MYC, p53^−/-^), SOAT1 knockout did not affect *in vitro* proliferation but significantly enhanced sensitivity to antigen-specific T cell killing (**Figures S1F-S1H**).

SOAT1 is an ER-resident multi-transmembrane enzyme that catalyzes cholesterol esterification for storage^22^. To investigate whether SOAT1’s enzyme activity contributes to cancer evasion of T cell-mediated killing, we re-expressed either wildtype SOAT1 or a catalytically inactive H460N mutant^19^ in SOAT1-knockout (SOAT1-KO) Hepa-OVA-Cas9 cells. When cocultured with antigen-specific OT-I CD8^+^ T cells, only wildtype SOAT1, not the H460N mutant, restored the resistance of SOAT1-KO cells to T cell killing (**Figures 1H and 1I**). These results highlight the importance of SOAT1’s enzymatic function in cancer-intrinsic evasion of T cell-mediated killing. To elucidate SOAT1’s role in tumorigenesis and immune evasion *in vivo*, we transplanted SOAT1-KO and vector-transduced Hepa1-6 cells into both immunodeficient Rag2^−/-^ mice and immunocompetent C57BL/6 mice. SOAT1 knockout had minimal effect on tumor growth in Rag2^−/-^ mice but significantly reduced tumor growth in C57BL/6 mice (**Figures 1J and 1K**). These findings suggest that SOAT1 is a critical driver of immune evasion in liver cancer.

### SOAT1 expression correlates with increased liver cancer recurrence and reduced overall survival

After confirming the role of SOAT1 in cancer evasion of T cell killing, we explored its clinical relevance in HCC prognosis and recurrence. Immunofluorescence staining of clinical samples showed that SOAT1 levels were significantly higher in HCC tissues compared to adjacent normal regions (**Figure S2A**). We then analyzed the correlation between SOAT1 expression and post-transplantation recurrence by measuring SOAT1 levels in primary HCC samples from patients with and without recurrence. The analysis revealed that primary HCCs from patients with recurrence had significantly higher SOAT1 levels than those from patients without recurrence (**Figure S2B**). Additionally, we examined SOAT1 levels in postsurgical recurred HCC samples, comparing those with secondary recurrence after liver transplantation to those without recurrence. Postsurgical recurred HCC samples from patients with secondary recurrence exhibited significantly higher SOAT1 levels than those without secondary recurrence (**Figure S2C and S2D**).

To assess whether SOAT1 expression could serve as an independent prognostic marker, we stratified liver transplant patients with HCC into two groups: SOAT1-high and SOAT1-low, using the average intratumoral SOAT1 score as the cutoff (**Figure 1L**). Patients with high intratumoral SOAT1 expression exhibited significantly higher rates of cancer recurrence and worse overall survival following liver transplantation compared to those with low SOAT1 expression (**Figures 1L and 1M**). In a subgroup analysis of patients receiving liver transplantation for primary HCC or postsurgical recurred HCC, high intra-tumoral SOAT1 levels were consistently associated with worse overall survival in both cohorts (**Figures S2E and S2F**). Moreover, high peri-tumoral SOAT1 levels were also correlated with significantly worse overall survival (**Figure 1N**). These findings suggest that high SOAT1 expression is a potential risk factor for HCC recurrence and is associated with poor overall survival.

### SOAT1 knockout or inhibition sensitizes cancer cell to T cell-mediated killing *in vitro* and *in vivo*

To determine whether SOAT1 knockout sensitizes tumors to T cell killing *in vivo*, we transplanted SOAT1-KO and vector control Hepa-OVA-Cas9 cells into Rag2^−/-^ mice, followed by OT-I CD8^+^ T cell therapy. Vector control tumors were resistant to T cell therapy, but SOAT1-KO tumors exhibited significantly reduced tumor growth after T cell treatment (**Figure 2A**). Next, we explored the role of SOAT1 in anti-PD1 immunotherapy by transplanting SOAT1-KO and vector control Hepa1-6 cells into C57BL/6 mice, followed by anti-PD1 treatment. SOAT1-KO tumors responded dramatically to anti-PD1 therapy, whereas vector control tumors did not (**Figure 2B**). Additionally, treatment with the SOAT1 chemical inhibitor avasimibe significantly enhanced the sensitivity of Hepa1-6 tumors to anti-PD1 therapy, similar to the effects of SOAT1 knockout (**Figure 2C**).

**Figure 2.**
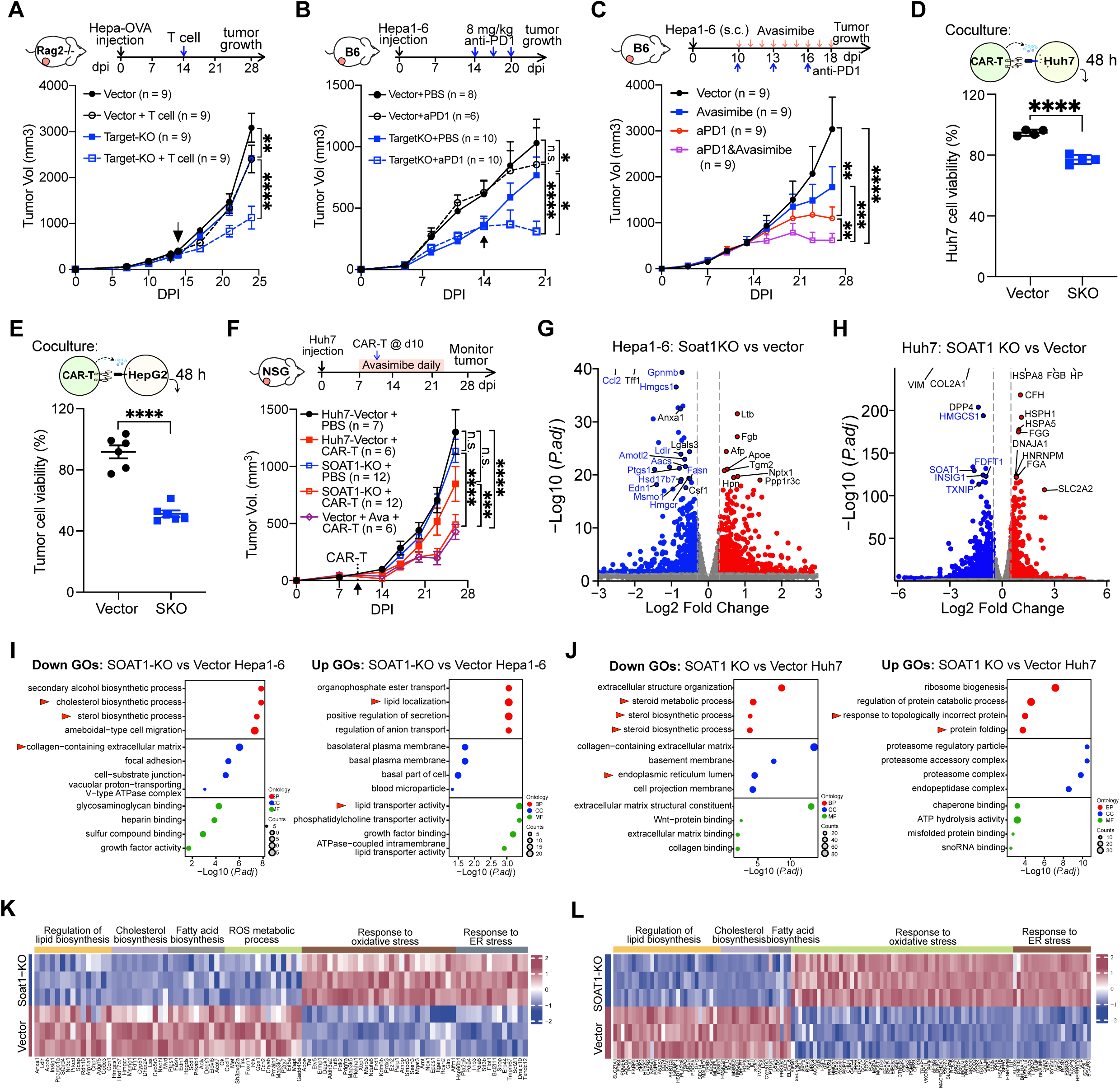
SOAT1 knockout enhances sensitivity of liver cancer cells to T cell-based immunotherapy. (**A**) Schematic of T cell therapy (upper panel). Growth curves of SOAT1-KO and vector control Hepa-OVA-Cas9 tumors in Rag2^−/-^ mice treated with PBS or OT-I CD8+T cells on day 14 post-tumor inoculation (n = 9 per group). (**B**) Schematic of anti-PD1 therapy (upper panel). Growth curves of SOAT1-KO (n = 10) and vector Hepa1-6 tumors in C57BL/6 mice (n = 8 for PBS, n = 6 for anti-PD1), treated with PBS or anti-PD1 starting on day 14, with 3 doses administered every 3 days. (**C**) Schematic of avasimibe and anti-PD1 combination therapy (upper panel). Growth curves of Hepa1-6 tumors in C57BL/6 mice treated with vehicle, avasimibe, anti-PD1, or their combination. Avasimibe was administered daily for 8 doses, and anti-PD1 every 3 days for 3 doses, starting on day 10. (**D, E**) Viability of SOAT1-KO and vector control Huh7 cells (**H**) and HepG2 cells (**I**) after 48 hours of coculture with GPC3-specific CAR-T cells at an E: T = 1: 1. (n = 4 per group). (**F**) Top: Schematics of CAR-T therapy combined with SOAT1 inhibition. Bottom: Tumor growth curves of SOAT1-KO and vector control Huh7 tumors in NSG mice treated with PBS, CAR-T cells, or a combination of avasimibe (daily starting on day 10) and CAR-T cells (administered once at day 10). (**G, H**) Volcano plots depicting differentially expressed genes between SOAT1-KO and vector control Hepa1-6 cells (**G**), and Huh7 cells (**H**). (**I, J**) GO analysis showing downregulated (left) and upregulated (right) processes in SOAT1-KO versus vector control Hepa1-6 cells (**I**) and Huh7 cells (**J**). (**K, L**) Heatmaps illustrating the differential expression of genes related to cholesterol biosynthesis, fatty acid biosynthesis, regulation of lipid biosynthesis, response to oxidative stress, and ER stress response in SOAT1-KO versus vector control Hepa1-6 cells (**K**) and Huh7 cells (**L**). Data are shown as means ± SEMs; Statistical analysis was performed using a Student’s t test or two-way ANOVA. Asterisks: * P < 0.05, ** P < 0.01, *** P < 0.001, ****P <0.0001. See also: Supplementary Figures S3.

We investigated whether SOAT1 knockout sensitizes human liver cancer cells to T cell-mediated killing. After confirming SOAT1 knockout in human liver cancer cell lines, Huh7 and HepG2 (**Figures S3A and S3B**), we assessed its effect on cancer cell proliferation. CCK-8 assays indicated that SOAT1 knockout had minimal impact on proliferation (**Figures S3C and S3D**). We then generated glypican-3 (GPC3)-specific chimeric antigen receptor (CAR)-T cells to evaluate the impact of SOAT1 knockout on cancer cell sensitivity to CAR-T therapy. Coculture experiments revealed that SOAT1-KO Huh7 and HepG2 cells were more sensitive to CAR-T cell-mediated killing (**Figures 2D and 2E**). To confirm this *in vivo*, we transplanted SOAT1-KO and vector control Huh7 cells into NSG mice, followed by GPC3 CAR-T cell therapy 10 days later. SOAT1 knockout significantly increased the sensitivity of human liver tumors to CAR-T therapy, without affecting their growth in immunodeficient NSG mice (**Figures 2F and S3E**). Notably, treatment with the SOAT1 inhibitor avasimibe similarly enhanced the therapeutic efficacy of CAR-T against liver cancer (**Figure 2F**). These results demonstrate that SOAT1 knockout enhances the responsiveness of human liver cancers to T cell-based therapies without significantly impacting tumor proliferation.

Next, we aimed to elucidate how SOAT1 knockout influences sensitivity to T cell therapy. We performed bulk RNA-seq analysis on SOAT1-KO and vector-transduced Hepa1-6 cells (**Figure 2G**) and Huh7 cells (**Figure 2H**). GO analysis of SOAT1-KO versus vector Hepa1-6 cells revealed significant downregulation of pathways related to cholesterol biosynthesis, sterol biosynthesis, collagen—containing extracellular matrix, and focal adhesion, while pathways associated with lipid transporter activity, phosphatidylcholine transporter activity, and lipid location were upregulated (**Figures 2I**). GSEA analysis further confirmed the downregulation of cholesterol, lipid metabolism, and prostaglandin synthesis and regulation (**Figures S3E-S3G**). Similarly, GO analysis of SOAT1-KO versus vector Huh7 cells showed downregulation of processes related to extracellular structure organization, steroid metabolism and biosynthesis, and collagen-containing extracellular matrix, with upregulation of processes linked to ribosome biogenesis, response to topologically incorrected protein, proteasome complex, and chaperone binding (**Figure 2J**). Further KEGG and metabolic GO analyses confirmed that SOAT1 knockout downregulated pathways involved in steroid biosynthesis, glycerolipid metabolism, and ECM-receptor interactions, while upregulating pathways for proteasome, complement and coagulation cascades, fatty acid degradation, and ER (endoplasmic reticulum) stress response (**Figures S3H and S3I**). Collectively, in both mouse and human cancer cells, we observed downregulation of genes associated with cholesterol biosynthesis, fatty acid biosynthesis, as well as reactive oxygen species (ROS) metabolism, while genes involved in response to oxidative stress and ER stress were upregulated in SOAT1 knockout cells (**Figures 2K and 2L**). These findings suggest that SOAT1 knockout enhances cancer cell sensitivity to T cell killing, likely by impairing their ability to cope with oxidative stress and ER stress.

### SOAT1 knockout modulates cancer transcriptome in vivo, reshaping the immune microenvironment

To explore how SOAT1 knockout sensitizes tumors to T cell-based immunotherapy *in vivo*, we analyzed the tumor immune microenvironment fostered by SOAT1-KO and vector-transduced Hepa1-6 cells in C57BL/6 mice treated with PBS or anti-PD1 (**Figure S4A**). SOAT1-KO tumors exhibited significantly higher percentages of tumor-infiltrating CD8^+^ T cells and CD4^+^ T cells, and lower levels of infiltrating monocytes compared to vector controls, in both PBS- and anti-PD1-treated groups (**Figures 3A-3C**). Besides, SOAT1-KO tumors displayed slightly higher infiltrating dendritic cells (DCs), particularly after anti-PD1 therapy (**Figure 3D**). The proportions of tumor-infiltrating B cells, macrophages and neutrophils were not significantly altered by SOAT1 knockout (**Figures S4B-S4D**). Notably, SOAT1-KO tumors showed an increase in PD1^+^CD8^+^ T cells, an effect further enhanced by anti-PD1 therapy (**Figures 3E and S4E**), suggesting a higher abundance of tumor-reactive CD8^+^ T cells in SOAT1-KO tumors. Collectively, these findings indicate that SOAT1 knockout sensitizes tumors to immunotherapy by promoting tumor-reactive T cell infiltration.

**Figure 3.**
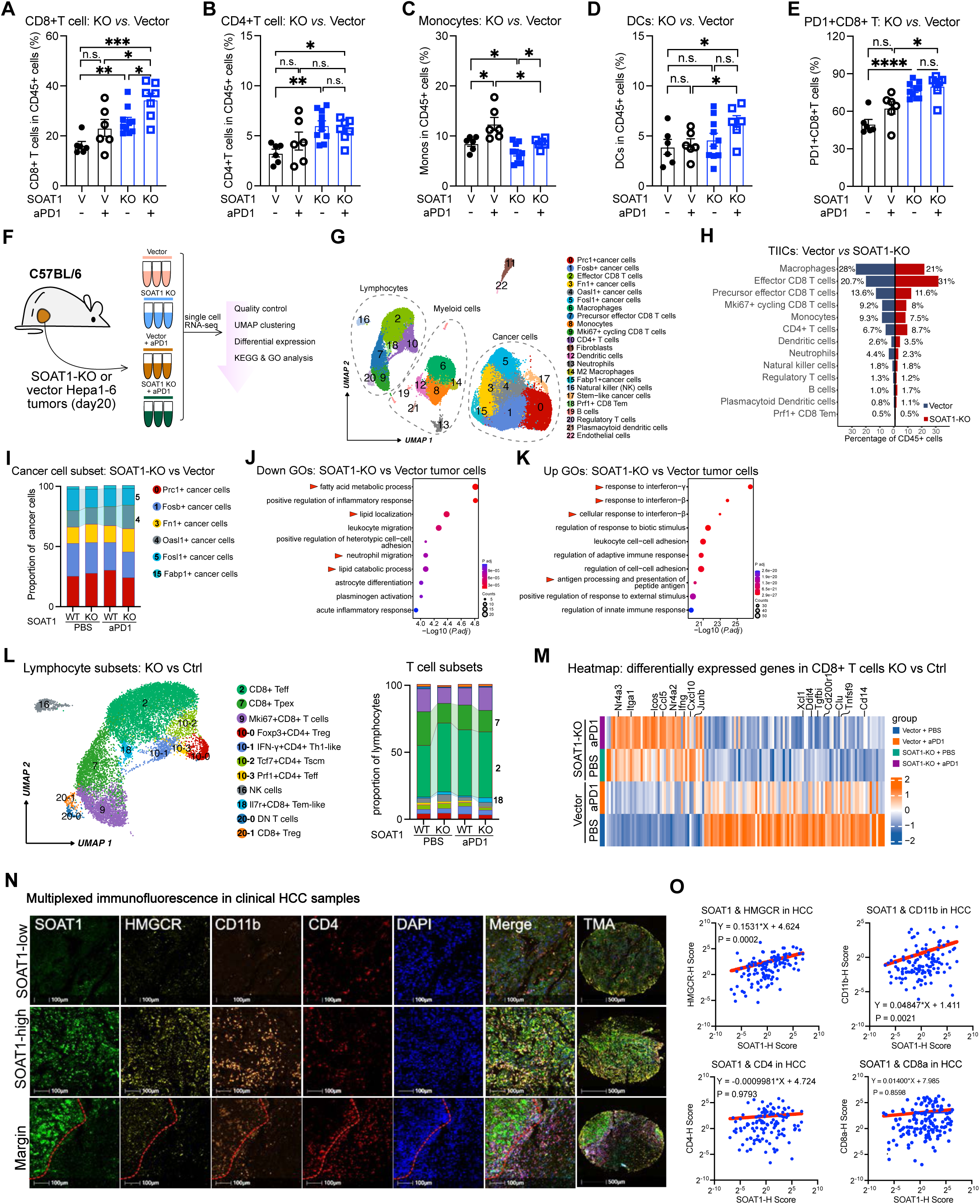
SOAT1 knockout reshapes tumor immune microenvironment with enhanced CD8+ T cell infiltration and effector function. (**A-E**) Flow cytometry analysis of the tumor immune microenvironment: quantification of CD8+ T cells (**A**), and CD4^+^ T cells (**B**), monocytes (**C**), and dendritic cells (**D**) among tumor-infiltrating CD45^+^ immune cells, and PD1^+^CD8^+^ T cells (**E**) in SOAT1-KO (n = 7) or vector control Hepa1-6 tumors (n = 6) from mice treated with PBS or anti-PD1. (**F**) Schematic illustrating single-cell RNA-sequencing (scRNA-seq) analysis of the immune microenvironment of SOAT1-KO and vector control Hepa1-6 tumors from C57BL/6 mice treated with PBS or anti-PD1 (pooled from 3 mice per group). (**G**) A uniform manifold approximation and projection (UMAP) plot showing 23 clusters identified by integrated analysis, with cell types annotated by cluster. (**H**) Bar plot showing proportional differences in immune cell subsets among tumor-infiltrating CD45+ immune cells in SOAT1-KO tumors compared to vector control tumors in C57BL/6 mice treated with PBS. (**I**) Bar plot depicting proportional differences in cancer cell subsets from SOAT1-KO and vector control Hepa1-6 tumors in mice treated with PBS or anti-PD1. (**J-K**) GO analysis showing downregulated (**J**) and upregulated (**K**) pathways in SOAT1-KO Hepa1-6 tumors compared to vector control tumors. (**L**) UMAP plot of scRNA-seq profiles showing 11 clusters among tumor-infiltrating T cells (left). Barplot depicting proportional differences in lymphocyte subsets between SOAT1-KO and vector control Hepa1-6 tumors in mice treated with PBS or anti-PD1 (right). (**M**) Heatmap of significant differential genes between CD8+T cells from SOAT1-KO versus vector Hepa1-6 tumors in mouse treated with PBS or anti-PD1. (**N, O**) Multiplexed immunofluorescence staining for SOAT1, HMGCR, CD11b, and CD4 in human HCC TMAs, assessing the correlation between SOAT1 expression, cholesterol biosynthesis, CD11b^+^ myeloid cells, and CD4^+^ T cells. Representative immunofluorescence images showing HCC samples with low, high, or hybrid SOAT1 expression (**N**), and correlation plots between SOAT1 expression and HMGCR, CD11b, CD4, and CD8a (**O**). Data are shown as means ± SEMs; Statistical analysis was performed using a Student’s t test or two-way ANOVA. Asterisks: * P < 0.05, ** P < 0.01, *** P < 0.001, **** P < 0.0001. See also: Supplementary Figures S4, S5, and S6.

To investigate the mechanism, we performed single-cell RNA sequencing (scRNA-seq) to profile the *in vivo* transcriptome and immune landscape (**Figure 3F**). Tumors from PBS- and anti-PD1-treated mice were purified using Ficoll and subjected to scRNA-seq, recovering 9,431 and 9,970 cells from SOAT1-KO tumors, and 10,880 and 8,580 cells from vector tumors, respectively. Unsupervised clustering identified 23 distinct cell clusters, with identities assigned based on differentially expressed genes (**Figure 3G**). In the PBS groups, SOAT1-KO tumors showed an increase in effector CD8^+^ T cells, CD4^+^ T cells, and DCs, while macrophages, monocytes, and neutrophils were reduced compared to vector tumors. The proportions of precursor effector CD8^+^ T cells were slightly lower in SOAT1-KO tumors, likely due to their transition to effector CD8^+^ T cells (**Figure 3H**). Interestingly, anti-PD1 treatment increased the percentage of effector CD8^+^ T cells and slightly reduced precursor effector CD8^+^ T cells in vector tumors (**Figure S5A**), similar to the effect of SOAT1 knockout. In the anti-PD1-treated groups, SOAT1-KO tumors had relatively higher proportions of DCs compared to vector controls (**Figure S5A**). These findings suggest that SOAT1 knockout enhances the infiltration of CD8^+^ T cell and DCs into tumors.

Analysis of cancer cell clusters identified six distinct subsets. SOAT1 knockout reduced the *Fosl1*+ cancer cell subset while increasing the interferon-relevant *Oasl1*^+^ cancer cell subset (**Figures 3I and S5B**). GO analysis showed that processes related to fatty acid metabolic process, lipid localization, and lipid catabolic process were downregulated in SOAT1-KO cancer cells, while response to IFN-γ and FN-β and antigen processing and presentation of peptide antigen were upregulated (**Figures 3J, 3K, S5C, and S5D**), consistent with bulk RNA-seq data that SOAT1 knockout suppresses cholesterol and lipid metabolism.

To assess how SOAT1 knockout affects tumor-infiltrating immune cells, we analyzed myeloid cell subsets. SOAT1-KO tumors exhibited higher percentages of DC subsets (cDC1, cDC2, and migratory DCs) and TCR+ macrophage, but lower percentages of *Spp1*^+^ macrophage and neutrophils (**Figures S5E and S5F**). GO analysis of myeloid transcriptomes revealed downregulation of processes related to the ERK1/ERK2 cascade, platelet alpha/secretory granule, and collagen-containing extracellular matrix, while processes involved in response to IFN-γ, antigen processing and presentation of exogenous antigen, and MHC-II complex were upregulated (**Figures S5G**). KEGG analysis demonstrated the downregulation of cholesterol metabolism and ferroptosis pathways, along with upregulation of phagosome and antigen processing and presentation pathways in the myeloid cells from SOAT1-KO tumors (**Figure S5H**). Furthermore, SOAT1-KO tumors exhibited higher percentages of NK cells, effector CD8^+^ T cells and Il7r^+^CD8^+^ effector memory like T cells (Tem) compared to vector controls (**Figures 3L and S5I**), with CD8^+^ T cells from SOAT1-KO tumors expressing higher levels of genes related to activation and recruitment (**Figure 3M**). GO analysis of CD8^+^ T cell transcriptomes indicated downregulation of plasminogen activation and negative regulation of extrinsic apoptotic signaling pathway via death domain receptors, but upregulation of chemokine/cytokine receptor binding pathways (**Figure S5J**). These results suggest that SOAT1 knockout disrupts lipid and cholesterol metabolism in both cancer cells and immune cells, while enhancing antigen presentation and immune responses.

To explore the relationship between SOAT1 expression, cholesterol biosynthesis, and immune cell infiltration in clinical samples, we performed multiplexed immunofluorescence (IF) staining on human tissue arrays (TMAs) of HCC and adjacent normal liver tissues. Our analysis revealed a positive correlation between the expression of HMGCR, a key enzyme in cholesterol biosynthesis, and SOAT1 expression in both HCC and adjacent normal tissues (**Figures 3N, 3O, S6A, and S6B**), highlighting the link between cholesterol biosynthesis and cholesteryl ester formation. Furthermore, we found that SOAT1 expression was positively correlated with CD11b expression in HCC but not in normal tissues, suggesting that elevated cholesteryl ester metabolism may be linked to the recruitment of CD11b^+^ myeloid cells. Intriguingly, we noted colocalization of SOAT1 with HMGCR and CD11b, along with some degree of colocalization between CD11b and CD4 at tumor-stroma margin (**Figures 3N, 3O, and S6C**). However, no significant correlation was found between SOAT1 expression and the infiltration of CD4^+^ or CD8^+^ T cells (**Figures 3N, 3O, S6B and S6D**). These findings suggest that SOAT1 expression is positively correlated with cholesterol metabolism and CD11b^+^ myeloid cell infiltration in HCC samples, indicating its potential role in modulating tumor microenvironment.

### SOAT1 knockout disrupts intratumoral cholesterol and lipid metabolism

Given that both *in vitro* RNA-seq and *in vivo* scRNA-seq data indicated downregulation of cholesterol and fatty acid biosynthesis genes in SOAT1-KO cells, we measured total free cholesterol and accessible plasma membrane cholesterol levels using Filipin III and Alexa Fluor 647-conjugated ALOD4 (AF647-ALOD4) staining, respectively. Interestingly, SOAT1-KO Hepa1-6 cells exhibited slightly increased total free cholesterol compared to vector controls, likely due to the cessation of cholesterol esterification (**Figure S7A**), as SOAT1 overexpression reduced total free cholesterol levels (**Figure S7B**). However, accessible plasma cholesterol was lower in SOAT1-KO Hepa1-6 cells (**Figure 4A**). Filipin III staining images showed that SOAT1 knockout altered cholesterol localization, sequestering more cholesterol in intracellular organelles rather than in the plasma membrane (**Figure S7C**). To assess whether this effect was consistent across different cell lines, we measured total free and accessible plasma membrane cholesterol in HCC51 (mouse) and Huh7 and HepG2 (human) liver cancer lines. SOAT1 knockout decreased either total free or/and accessible plasma membrane cholesterol in these cells (**Figures 4B and S7D-S7F**).

**Figure 4.**
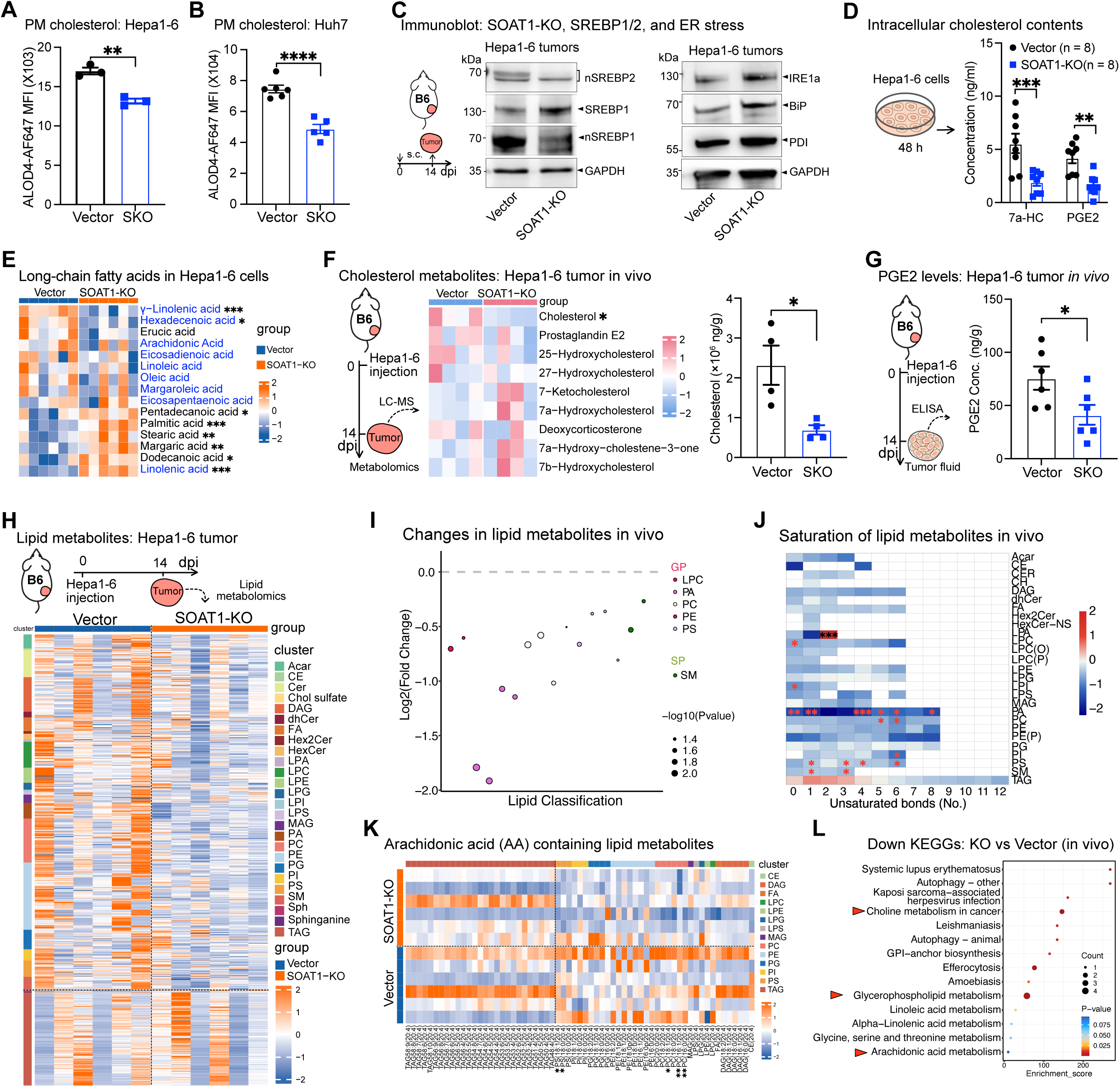
SOAT1 knockout inhibits intratumoral cholesterol biosynthesis and lipid metabolism. (**A-B**) Accessible plasma membrane cholesterol in SOAT1-KO and vector control Hepa1-6 cells (**A**) (n = 3) and Huh7 cells (**B**) (n ≥ 5) was assessed by ALOD4-AF647 staining and analyzed by flow cytometry. (**C**) Immunoblot assays showing reduced processing of SREBP2 into nSREBP2, SREBP1 into nSREBP1, and activation of ER stress proteins (IRE1a, BiP, PDI) were confirmed in SOAT1-KO versus vector control Hepa1-6 tumors collected from C57BL/6 mice at day 14 post-inoculation. (**D**) Bar plot shows intracellular 7a-HC and PGE2 levels in SOAT1-KO and vector Hepa1-6 cells based on cholesterol-targeted metabolomics analysis (n = 8). (**E**) Heatmap of intracellular long-chain fatty acids in SOAT1-KO versus vector Hepa1-6 cells, with unsaturated fatty acids labelled in blue (n = 6). (**F**) Cholesterol-targeted metabolomics on SOAT1-KO versus vector Hepa1-6 tumors. The heatmap (left) highlights intratumoral cholesterol metabolites, and bar plot (right) shows differences in intratumoral cholesterol levels (n = 4). (**G**) Intratumoral PGE2 levels in SOAT1-KO versus vector Hepa1-6 tumors, measured by ELISA (n = 6). (**H-K**) Lipid-targeted metabolomics of SOAT1-KO versus vector Hepa1-6 tumors (n = 6): (**H**) Heatmap showing relative abundance of lipid metabolites, with fold-changes in SOAT1-KO tumors compared to vector controls; (**I**) Bubble plot illustrating significant decreases in lipid classes, particularly glycerol phospholipids such as LPC, PA, PC, PE, PS; (**J**) Heatmap showing changes in lipid metabolites with varying degrees of unsaturation; (**K**) Heatmap of arachidonic acid-containing lipids, showing reduced phospholipids but unchanged TAG levels. (**L**) Downregulated KEGG pathways in SOAT1-KO tumors, based on differential lipid metabolite analysis. CE = cholesteryl esters, DAG = diacylglycerol, FA = fatty acids, LPC = lysophosphatidylcholine, LPE = lysophosphatidyl-ethanolamine, LPG = lysophosphatidylglycerol, LPS = lysophosphatidylserine, MAG = monoacylglycerol, PC = phosphatidylcholine, PE = phosphatidylethanolamine, PG = phosphatidylglycerol, PI = phosphatidylinositol, PS = phosphatidylserine, TAG = triacylglycerol. Data are shown as means ± SEMs; Statistical analysis was performed using a Student’s t test or two-way ANOVA. Asterisks: * P < 0.05, ** P < 0.01, *** P < 0.001, **** P < 0.0001. See also: Supplementary Figure S7

Considering SOAT1’s role in converting cholesterol into cholesteryl esters in the ER, we hypothesized that free cholesterol accumulation in SOAT1-KO cells inhibits SREBP1 and SREBP2 processing via SCAP binding, suppressing cholesterol and fatty acid biosynthesis. Immunoblotting confirmed lower levels of processed nSREBP1 and nSREBP2 in SOAT1-KO tumors from C57BL/6 mice, despite higher levels of unprocessed SREBP1 (**Figure 4C**). Additionally, ER stress proteins IRE1a, BiP, and PDI were elevated in SOAT1-KO tumors, in consistent with the scRNA-seq data (**Figure S7G**). These data indicate that SOAT1 knockout disrupts cholesterol and fatty acid biosynthesis by preventing SREBP processing.

### Lipidomic analysis profiled metabolic changes induced by SOAT1 knockout

Next, we performed targeted lipidomic and cholesterol metabolomic analyses on both SOAT1-KO and vector Hepa1-6 cells. The results demonstrated that SOAT1 knockout significantly reduced intracellular 7α-hydroxycholesterol (7α-HC) and prostaglandin E2 (PGE2) levels (**Figures 4D and S7H**). Lipidomics revealed a decrease in cholesteryl esters alongside an increase in intracellular triacylglycerol (TAG) levels *in vitro* (**Figures S7I and S7J**). Moreover, SOAT1-KO cells had elevated levels of saturated fatty acids but reduced levels of unsaturated fatty acids (**Figure 4E**), suggesting an alternation in fatty acid metabolism.

In vivo metabolomics on SOAT1-KO and vector Hepa1-6 tumors from C57BL/6 mice revealed significant lower intratumoral cholesterol and PGE2 levels, which was confirmed by ELISA analysis of tumor lysates (**Figures 4F, 4G, and S7K**). Lipidomics revealed that SOAT1-KO tumors had reduced levels of lipid metabolites, except for triacylglycerols (**Figure 4H**). Further analysis revealed significant reductions in glycerophospholipids, especially those containing unsaturated fatty acids in SOAT1-KO tumors (**Figures 4I, 4J, S7L, and S7M**). Reduced arachidonic acid (AA)-containing glycerophospholipids, but not TAGs, may account for the lower PGE2 production in SOAT1-KO tumors (**Figure 4K**). KEGG analysis indicated a decrease in pathways related to autophagy, glycerophospholipid metabolism, and unsaturated fatty acid (e.g. AA) metabolism in SOAT1-KO tumors (**Figure 4L**), potentially explaining reduced PGE2 levels.

### SOAT1 knockout sensitizes cancer cells to T cell killing by impairing anti-oxidant capacity

Due to the known immunosuppressive effects of PGE2^24,25^, we measured PGE2 secretion by SOAT1-KO and vector Hepa1-6 cells in the presence of T cell killing. SOAT1-KO cells released significantly less PGE2 under T cell-mediated killing, an effect reversed by re-expressing SOAT1 (**Figure 5A**). Supplementing PGE2 during T cell activation and cancer cell-T cell cocultures demonstrated that PGE2 suppressed CD8^+^ T cell proliferation and effector function in a dose-dependent manner (**Figures 5B and S8A**), highlighting the contribution of reduced PGE2 in increased T cell sensitivity.

**Figure 5.**
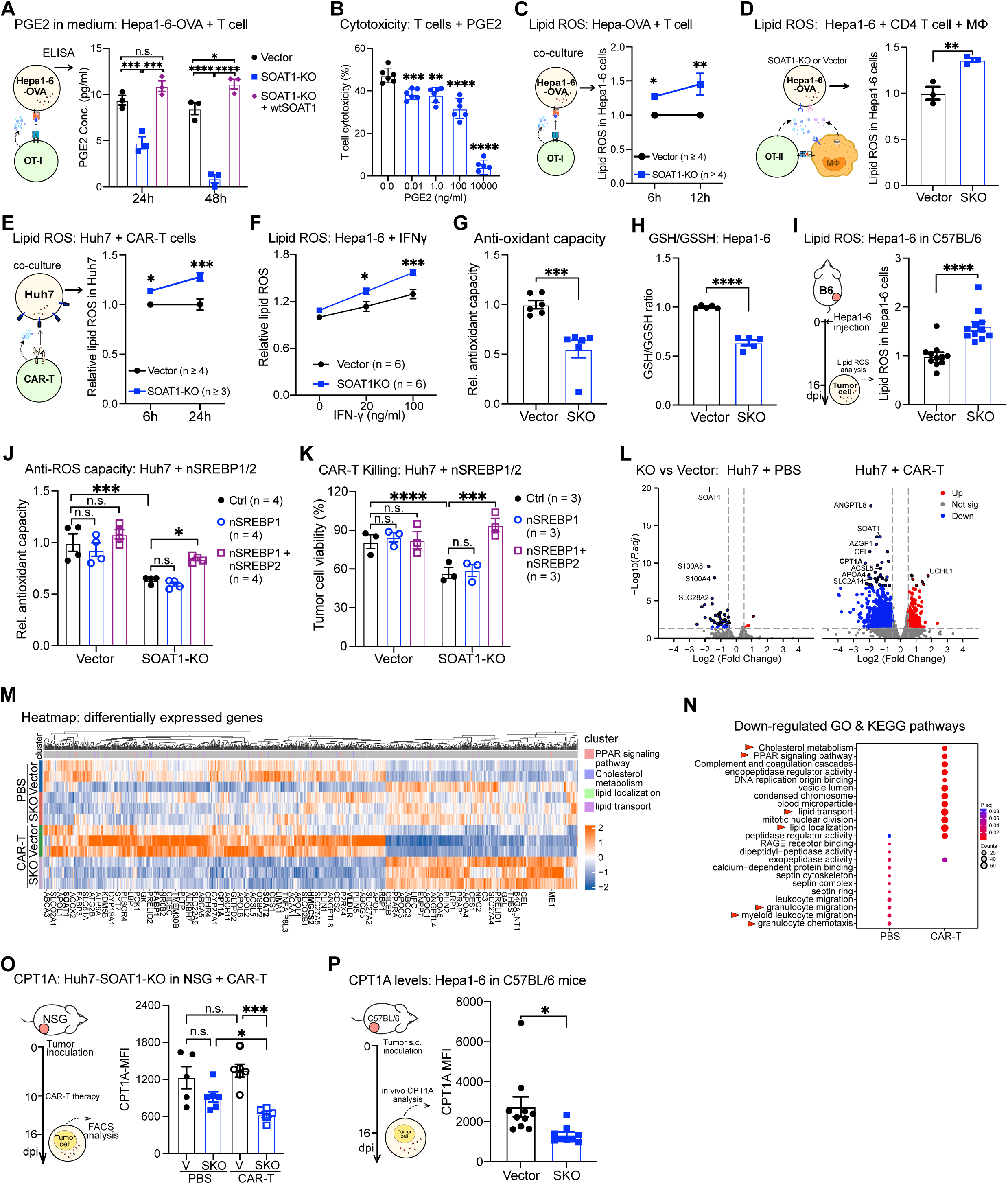
SOAT1 knockout senstizes cancer cell to T cell immunity by reducing PGE2 secretion and lowering antixodiant capacity. (**A**) The concentration of PGE2 in the medium of SOAT1-KO, vector, and wtSOAT1-rescued Hepa-OVA-Cas9 cells after coculture with antigen-specific OT-I CD8+ T cells was measured by ELISA (n = 3). (**B**) The impact of PGE2 on effector function of CD8+ T cells were assessed by adding PGE2 to the coculture of OT-I T cell with Hepa-OVA-Cas9 cells (n = 6). (**C, D**) Lipid ROS levels in SOAT1-KO versus vector Hepa-OVA-Cas9 cells were measured using the lipid peroxidation sensor BODIPY-C11 after coculture with OT-I CD8+ T cells (n ≥ 4) (**C**), and after coculture with bone-marrow-derived macrophage (BMDM) and OT-II CD4+ T cells at a ratio of 1:1:2 for 48 hours (n = 3) (**D**). (**E**) Lipid ROS levels in SOAT1-KO and vector Huh7 cells after coculture with GPC3-targeting CAR-T cells were measured (n = 5 at 6 h, n = 4 at 12 h). (**F**) Lipid ROS levels in SOAT1-KO and vector Hepa1-6 cells treated with vehicle or IFN-γ (n = 6). (**G, H**) The antioxidant capacity and oxidative status of SOAT1-KO and vector Hepa1-6 cells was evaluated using the Trolox Equivalent Antioxidant Capacity (TEAC) assay (n = 6) (**G**) and a GSH/GSSG ratio detection kit (n = 5) (**H**). (**I**) Lipid ROS levels *in vivo* were evaluated in SOAT1-KO and vector Hepa1-6 tumor cells from immunocompetent C57BL/6 mice via BODIPY-C11 staining (n = 11). (**J**) The anti-oxidant capacity of control, nSREBP1, or nSREBP1 and nSREBP2-overexpressed SOAT1-KO and vector Huh7 cells (n = 4). (**K**) Viability of control, nSREBP1, or nSREBP1 and nSREBP2-overexpressed SOAT1-KO and vector Huh7 cells when cocultured with CAR-T cells for 48 h (n = 3). (**L**) Volcano plots showing differentially expressed genes in SOAT1-KO versus vector control Huh7 tumors from NSG mice treated with PBS (left) and GPC3 CAR-T cells (right). (**M**) Heatmap showing the differentially expressed lipid metabolites among SOAT1-KO versus vector control Huh7 tumors from NSG mice treated by PBS or GPC3 CAR-T cells (n = 3). (**N**) GO & KEGG analysis showing significantly downregulated processes in SOAT1-KO Huh7 tumors compared to vector controls. (**O, P**) CPT1A expression was assessed in SOAT1-KO and vector Huh7 tumors from NSG mice treated with PBS or CAR-T cells (n = 5) (**O**), and in SOAT1-KO and vector control Hepa1-6 tumors from C57BL/6 mice by flow cytometry (n = 10) (**P**). Data are shown as means ± SEMs; Statistical analysis was performed using a Student’s t test or two-way ANOVA. Asterisks: * P < 0.05, ** P < 0.01, *** P < 0.001, **** P < 0.0001. See also: Supplementary Figure S8

Next, we examined whether SOAT1 knockout sensitizes cancer cells to perforin-mediated cell death. We used streptolysin O (SLO) to permeabilize both SOAT1-KO and control Hepa1-6 cells, and found no significant difference in their sensitivity to SLO-mediated permeabilization (**Figure S8B**). Given our RNA-seq data indicating increased oxidative and ER stress in SOAT1-KO cells, we hypothesized that SOAT1-KO may compromise cancer cells’ ability to manage oxidative stress triggered by T cell cytotoxicity. Indeed, under CD8^+^ T cell or CD4^+^ T cell plus macrophage-mediated killing, SOAT1-KO Hepa1-6 cells exhibited significantly higher lipid ROS levels than vector controls (**Figures 5C, 5D, and S8C**). This trend was also observed in SOAT1-KO Huh7 cells under GPC3 CAR-T cell killing (**Figures 5E and S8D**). Furthermore, direct treatment with IFN-γ resulted in increased lipid ROS levels in SOAT1-KO cells compared to controls (**Figure 5F**). In addition, the Trolox equivalent antioxidant capacity (TEAC) assay revealed that SOAT1-KO cells had significantly reduced antioxidant capacity (**Figure 5G**), while their glutathione (GSH) to oxidized glutathione (GSSG) ratios were also diminished (**Figure 5H**), confirming a loss of redox resilience. To explore whether the observed increase in lipid peroxidation occurs *in vivo*, we measured lipid ROS levels in cancer cells and tumor-infiltrating immune cells. SOAT1-KO cancer cells exhibited significantly higher lipid ROS levels than controls, though no difference was observed in the immune cells (**Figures 5I and S8E**). These findings suggest the reduced antioxidant capacity in SOAT1-KO cancer cells contributes to their increased sensitivity to T cell-mediated killing.

To further investigate the role of cholesterol and fatty acids biosynthesis in regulating antioxidant capacity and sensitivity to immune attack, we introduced the transcriptionally active form of nSREBP1, nSREBP2, or a combination of both into SOAT1-KO and vector Hepa1-6 cells and Huh7 cells. Notably, restoring nSREBP2 or both nSREBP2 and nSREBP1, but not nSREBP1, in SOAT1-KO Hepa1-6 cells and Huh7 cells significantly recovered their antioxidant capacity and restored the resistance to T cell-mediated killing (**Figures 5J, 5K, S8F, and S8G**). Furthermore, tumor growth curves in immunocompetent C57BL/6 mice revealed that re-expression of nSREBP2, but not nSREBP1, promoted tumor growth and eliminated the growth difference between SOAT1 knockout and vector control tumors (**Figures S8H and S8I**). These findings suggest that cholesterol biosynthesis plays a crucial role in maintaining redox resilience and facilitating immune evasion in cancer cells.

RNA-seq analysis of SOAT1-KO and vector Huh7 tumors treated with GPC3 CAR-T therapy revealed pronounced transcriptomic differences between SOAT1-KO and control tumors during T cell-mediated killing compared to PBS treatment (**Figures 5L and 5M**). GO and KEGG analyses indicated that SOAT1-KO tumors treated with PBS already displayed downregulation of pathways related to granulocyte and myeloid leukocyte migration (**Figure 5N**). In contrast, SOAT1-KO tumors under T cell therapy exhibited significant downregulation of pathways associated with cholesterol metabolism, PPAR signaling, lipid transport, and lipid localization (**Figures 5M and 5N**), consistent with our scRNA-seq (**Figure 3J**). PPAR-mediated fatty acid β-oxidation (FAO) has been indicated in cancer immune evasion^23,24^. Notably, we found that both SOAT1-KO human Huh7 tumors under CAR-T cell therapy and SOAT1-KO mouse Hepa1-6 tumors under immunosurveillance had significantly lower levels of carnitine palmitoyl-transferase 1A (CPT1A), an essential enzyme for transporting long-chain fatty acids into mitochondria for β-oxidation (**Figures 5O and 5P**). Overall, these results indicate that SOAT1 knockout suppresses cholesterol and fatty acid metabolism, leading to reduced anti-oxidant capacity and impaired redox and metabolic resilience under immune pressure.

### SOAT1 inhibition suppresses liver cancer occurrence and enhances immunotherapy efficacy under immunosuppressive conditions

Based on our finding that SOAT1 knockout significantly enhances the sensitivity of cancer cell to immune-mediated killing, we hypothesized that inhibiting SOAT1 could reduce cancer recurrence and improve the efficacy of immunotherapy, even in immunocompromised conditions. To test this, we developed a liver cancer mouse model with varying degrees of immunosuppression using immunosuppressants such as tacrolimus (a calcineurin inhibitor), sirolimus (an mTOR inhibitor), or a combination of both. As anticipated, immunosuppressant treatment markedly accelerated liver cancer development and growth, leading to significantly reduced overall survival (**Figures 6A, 6B, and S9A-S9C**). Next, we transplanted SOAT1-KO and vector Hepa1-6 cells into tacrolimus-treated mice to evaluate the effect of SOAT1 knockout on tumor occurrence and growth under compromised immunosurveillance. Remarkably, SOAT1 knockout significantly suppressed tumor occurrence and growth compared to vector controls, both in normal and immunocompromised conditions, but not in fully immunodeficient settings (**Figures 6C and S9D**). Furthermore, treatment with the SOAT1 inhibitor avasimibe significantly reduced liver tumor occurrence and growth in tracrolimus-induced immunocompromised models (**Figure 6D**). These findings demonstrate that targeting SOAT1, via genetic knockout or chemical inhibition, effectively reduces tumor occurrence and growth, even under suboptimal immune conditions.

**Figure 6.**
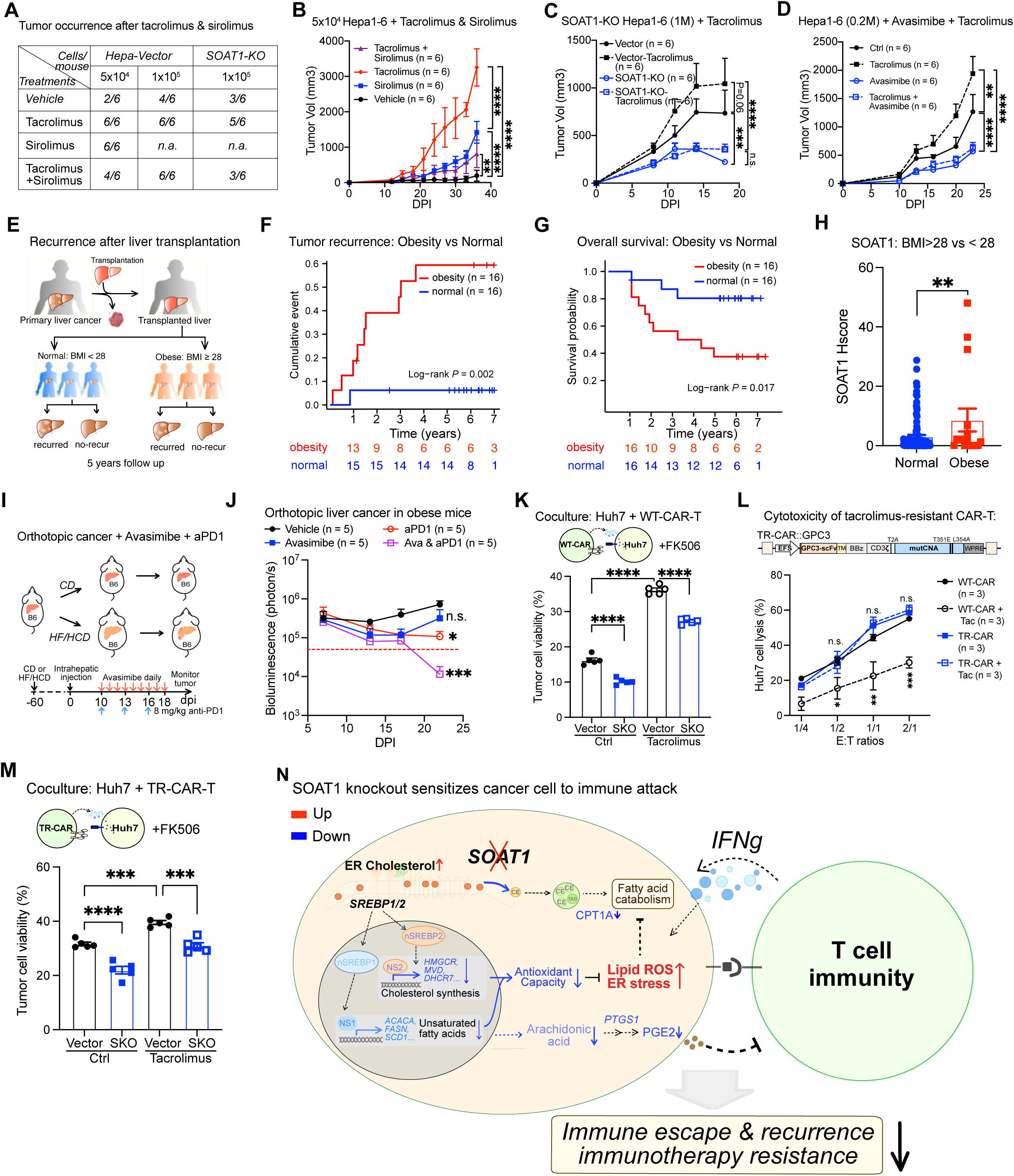
SOAT1 inhibition reduces cancer growth and enhances CAR-T therapy under suboptimal immunosurveillance. (**A**) Table presents the tumor occurrence rates in C57BL/6 mice transplanted with 5×10^4^ or 1×10^5^ SOAT1-KO or control Hepa1-6 cells, followed by treatment of PBS, sirolimus, tacrolimus, and their combination. (**B**) Growth curves of Hepa1-6 tumors in C57BL/6 mice subcutaneously inoculated with 5×10^4^ cells and treated with PBS, sirolimus, tacrolimus, and their combination (n = 6). (**C**) Tumor growth curves in C57BL/6 mice subcutaneously injected with 1×10^6^ SOAT1-KO or vector Hepa1-6 cells, and treated with PBS or Tacrolimus (n = 6). (**D**) Tumor growth curves in C57BL/6 mice subcutaneously inoculated with 2×10^5^ Hepa1-6 cells and treated with vehicle, tacrolimus, avasimibe, or their combination (n = 6). (**E**) Schematics depicting the study of HCC recurrence in patients stratified by body mass index (BMI). (**F, G**) The relationship between obesity and HCC recurrence (**F**) and overall survival (**G**) was analyzed by dividing the patients into two cohorts: BMI ≥ 28 (n = 16) and BMI < 28 (n = 16). (**H**) SOAT1 expression levels were compared between obese HCC patients and those with normal BMI. (**I**) Schematics depicting the experimental design using the SOAT1 inhibitor avasimibe and anti-PD1 treatment in orthotopic liver cancer in normal or obese C57BL/6 mice. Mice were fed with either a common diet (CD) or a high fat, high cholesterol diet (HF/HCD) to simulate different metabolic conditions. (**J**) Growth of orthotopic liver tumors in HF/HCD-fed obese mice, assessed by measuring luciferase activity. Mice were treated with vehicle, avasimbe, anti-PD1, or their combination starting on day 10 post-tumor inoculation. (n = 5) (**K**) Viability of SOAT1-KO and vector control Huh7 cells after coculture with WT-CAR-T cells at an E:T ratios of 1:1 in the presence of vehicle or tacrolimus (n = 4). (**L**) Upper panel: the design of tacrolimus resistant (TR)-CAR-T cells by expressing mutant calcineurin A (T351E, L354A). Lower panel: cytotoxic activity of WT-CAR-T cells and TR-CAR-T cells in the presence of tacrolimus or vehicle, measured by coculturing with Huh7 cells at various E:T ratios. Target cell lysis was quantified by the reduction of luciferase activity in Huh7 cells (n = 3). (**M**) Viability of SOAT1-KO and vector control Huh7 cells after coculture with TR-CAR-T cells at an E:T ratios of 1:1 in the presence of vehicle or tacrolimus (n = 4). (**N**) Graphical abstract illustrating how SOAT1 knockout reduces anti-oxidant capacity and increases cancer cell sensitivity to T cell-mediated immunity. Data are shown as means ± SEMs; Statistical analysis was performed using a Student’s t test or two-way ANOVA. Asterisks: * P < 0.05, ** P < 0.01, *** P < 0.001, **** P < 0.0001. **See also: Supplementary Figure S9 and Table S2.**

Given the association between obesity and compromised antitumor immunity^5,8^, we investigated whether SOAT1 contributes to obesity-associated HCC recurrence. Patients were categorized into normal (BMI < 28) and obese (BMI ≥ 28) groups based on body mass index (BMI) (**Table S2**). Obese patients exhibited significantly higher recurrence rates and worse overall survival compared to those with normal BMI (**Figures 6E-6G**). Since SOAT1 plays a critical role in converting excess cholesterol into cholesteryl esters stored in lipid droplets, we hypothesized that SOAT1 expression might be elevated in obese patients. Immunofluorescence staining confirmed that obese patients had significantly higher SOAT1 expression than those with normal BMI (**Figure 6H**), indicating a potential link between SOAT1 and obesity-associated HCC recurrence.

To investigate whether SOAT1 inhibition could suppress orthotopic liver cancer growth and enhance immunotherapy responses in obese conditions, we established an obesity-associated tumor model by feeding C57BL/6 mice a high fat, high cholesterol diet (HF/HCD) for 2 months, followed by orthotopic liver cancer transplantation (**Figure 6I**). In this model, orthotopic liver tumors in both normal and obese mice were profoundly suppressed by the combination of avasimibe and anti-PD1 therapy (**Figures 6J, S9E, and S9F**). These findings suggest that SOAT1 inhibition not only enhances immunotherapy under normal conditions, but also substantially boosts the antitumor efficacy of anti-PD1 immunotherapy in obese mice.

Additionally, we explored whether SOAT1 inhibition could enhance immunotherapy for relapsed HCC under immunosuppressive conditions, simulating the environment of liver transplant recipients. Intriguingly, SOAT1 knockout significantly increased the vulnerability of Huh7 cells to WT-CAR-T cell-mediated killing even in the presence of tacrolimus which severely inhibited WT-CAR-T cell cytotoxicity (**Figure 6K**), further corroborating that SOAT1 inhibition sensitizes cancer cells to even compromised immune surveillance. To enable more effective CAR-T therapies against recurred HCC in the presence of immunosuppressant, we engineered a tacrolimus-resistant CAR (TR-CAR) by incorporating a mutant form of calcineurin A (T351E and L354A) into the CAR construct^28^. Functional assays confirmed that TR-CAR-T cells retained their cytotoxic activity in the presence of tacrolimus (FK506), both in vitro and in vivo, whereas the original CAR-T cells were significantly impaired under these conditions (**Figure 6L**). To evaluate the potential synergy between SOAT1 inhibition and CAR-T therapy in the presence of immunosuppressants, we co-cultured SOAT1-KO and vector control Huh7 cells with both WT-CAR-T cells and TR-CAR-T cells, with tacrolimus added to the medium. Notably, SOAT1-KO cancer cells exhibited higher sensitivity to TR-CAR-T cell-mediated killing under both tacrolimus-treated and untreated conditions (**Figure 6M**). These results further support the idea that SOAT1 inhibition sensitizes cancer cells to immune attack, even when immune function is compromised by immunosuppressants.

Collectively, these findings demonstrate that SOAT1 inhibition reduces liver cancer occurrence and growth and significantly boosts CAR-T efficacy against liver cancer even under immunocompromised conditions.

## Discussion

In this study, we combined differential proteomics in recurrent versus non-recurrent from clinical cohorts with functional assays to identify key drivers of immune evasion and recurrence in liver cancer. Through this approach, we identified SOAT1 as a critical regulator of redox metabolic balance, promoting cancer-intrinsic immune evasion and recurrence. SOAT1 knockout in both mouse and human liver cancer cells significantly enhanced their sensitivity to host immunosurveillance and T cell-mediated immunotherapy. Bulk RNA-seq, scRNA-seq, and targeted metabolomics revealed that SOAT1 knockout reduces intratumoral cholesterol and lipid metabolism in tumors while enhancing their IFN-β / IFN-γ responses and antigen presentation. These metabolic alternations impair the tumors’ ability to manage oxidative stress and ER stress under T cell attack. Tumor microenvironment analysis via flow cytometry and scRNA-seq further demonstrated that SOAT1 knockout increases the infiltration of CD8^+^ T cells and dendritic cells while reducing neutrophil and M2 macrophage infiltration. Cholesterol and lipid metabolism are known to support tumor progression. Here, we demonstrate that SOAT1-mediated cholesterol esterification acts as a key regulator coordinating lipid metabolism to bolster the redox resilience of cancer cells, thereby driving immune evasion and resistance to immunotherapy in liver cancer.

High SOAT1 expression has been associated with progression across multiple cancer types^29,30^. Previous study has directly linked SOAT1 expression to cancer cell proliferation by *in vitro* shRNA knockdown and avasimibe inhibition experiments^30^. In contrast, we found that SOAT1 knockout in both mouse (Hepa1-6 and HCC51) and human (Huh7 and HepG2) liver cancer cell lines had minimal effect on their *in vitro* proliferation and *in vivo* tumorigenesis in immunodeficient mice. Rather, SOAT1 knockout dramatically sensitized both mouse and human liver cancer cells to host immunosurveillance, T cell killing, and anti-PD1 immunotherapy. Both in vitro and in vivo RNA-seq analysis showed that SOAT1 knockout inhibits endogenous cholesterol and fatty acid biosynthesis while elevating the oxidative and ER stress. Indeed, metabolomics showed that SOAT1 knockout significantly decreased intratumoral cholesterol, unsaturated fatty acids, phospholipids, and PGE2 in immunocompetent mice or under T cell attack. These data indicate that SOAT1-mediated cholesterol esterification plays a key role in immune evasion and immunotherapy resistance of liver cancer.

Further investigations revealed that SOAT1 knockout significantly impaired the antioxidant capacity of cancer cells, leading to elevated levels of lipid peroxidation under T cell killing or direct IFN-γ treatment. As SOAT1 converts excess cholesterol into cholesteryl esters in the ER, its knockout resulted in cholesterol accumulation in the ER, preventing the processing of SREBP1 and SREBP2 for cholesterol and fatty acid biosynthesis. Given the reported antioxidant roles of various cholesterol intermediates and unsaturated fatty acids^31,32^, we reason that SOAT1 knockout impairs the antioxidant capacity of liver cancer cells by disrupting these metabolic pathways. Moreover, *in vivo* studies showed that SOAT1 knockout reduces FAO catabolism, as evidence by lower CPT1A levels, and decreases PGE2 release through reduced arachidonic acid metabolism, particularly under immunosurveillance or T cell therapy stress. Collectively, SOAT1 knockout enhances cancer cell susceptibility to immune-mediated lipid peroxidation and promotes T cell effector functions by reducing the release of immunosuppressive metabolites like PGE2^24,25^. Unlike direct inhibition of enzymes in cholesterol or fatty acid biosynthesis, SOAT1 inhibition does not impair lipid biosynthesis essential for basal cellular functions, as indicated by the non-lethality and normal development of SOAT1 knockout mice^33^. Instead, SOAT1 inhibition abolishes cancer cells’ cholesterol metabolism advantages in utilizing excessive cholesterol biosynthesis to cope with oxidative stress imposed by immune surveillance and immunotherapy, thereby sensitizing cancer cells to T cell-mediated killing (**Figures 6N and S9G**).

Importantly, targeting SOAT1 via genetic knockout or pharmacological inhibitors improves the efficacy of T cell-based immunotherapies and reduces the growth and recurrence of liver tumors under both normal and compromised immunosurveillance conditions. For example, SOAT1 inhibition markedly reduced liver cancer occurrence and growth even in the presence of immunosuppressants, and boosts anti-PD1 therapy efficacy in an obese mice model. Liver transplantation is one of the most effective treatments for HCC, but immunosuppressants like calcineurin inhibitors are commonly used to prevent rejection and maintain graft function^34^. Since host immunosurveillance is critical for preventing cancer recurrence, immunosuppressant-induced immunosuppression has become an independent risk factor for post-transplant cancer recurrence. Our study suggests that SOAT1 inhibition has potential in preventing liver cancer recurrence even under immunosuppressant-mediated compromised immunosurveillance conditions. Besides, SOAT1 inhibition has been shown to enhance T cell antitumor functions , further supporting its potentials in boosting cancer immunotherapy^35^.

Our clinical sample analyses strongly supported SOAT1’s role in liver cancer immune evasion and recurrence. Multiplexed immunofluorescence staining of human HCC samples and follow-up data demonstrated SOAT1 expression correlates with active cholesterol metabolism and CD11b^+^ myeloid cell infiltration, as well as the increased HCC recurrence and worse overall survival. Additionally, our clinical data showed that SOAT1 expression is higher in obese patients, suggesting SOAT1 may contributes to obesity-associated liver cancer immune evasion and recurrence. Given the rising prevalence of obesity and associated metabolic disorders, the therapeutic potential of targeting SOAT1 warrants further investigation.

In summary, we employed a clinical data-driven functional screen to identify SOAT1 as a key metabolic driver of cancer-intrinsic immune evasion and recurrence. Mechanistic studies reveal that SOAT1 serves as a metabolic hub in lipid metabolism, enhancing the antioxidant capacity of cancer cells by facilitating excessive cholesterol biosynthesis through cholesterol esterification. Blocking cholesterol esterification via SOAT1 inhibition disrupts liver cancer’s advantages in leveraging robust cholesterol biosynthesis to mitigate oxidative stress, thereby sensitizing HCC to both immune surveillance and immunotherapy. Notably, SOAT1 expression is significantly elevated in obese patients. Given the increasing prevalence of obesity and hypercholesterolemia, along with their association with cancer risk, the role of cholesterol esterification in driving immune evasion across other cancer types warrants further investigation. Overall, our findings identify SOAT1-mediated cholesterol esterification as a critical mechanism underlying immune evasion and recurrence in liver cancer, positioning SOAT1 as a promising target to overcome immunotherapy resistance and prevent HCC recurrence.

## Acknowledgements

We thank Baojin Wu and Guoyuan Chen for animal husbandry and imaging, Wei Bian for the technical support on flow cytometry at the CEMCS Core Facility. This study is supported by the National Key Research and Development Program of China (2023YFC2505900), the Strategic Priority Research Program of the Chinese Academy of Science (Grant No. XDB0990000), National Natural Science Foundation of China (82241226, 32170917, 82241225, 81873874, 82071797), and Shanghai Original Exploratory Program (23ZR1481500). We thank Prof. Dan Li, Prof. Lu Lu, Prof. Quanbao Zhang, and all members from Guangchuan Wang Lab and Zhengxin Wang Lab for the helpful discussions and technical support at the Lab meetings.

## Author Contributions

G.W. conceived the project. G.W. Y.G. Z.W. and J. L. designed the experiments. Y.G. performed most of the experiments. L.Z. performed CRSIPR screening and human CAR-T therapy experiments with W.H. and Y.G.’s help. J. L. performed proteomics analysis of clinical samples. W.H., X.D., Y.L., Y.H., Z.N., L.B., Y.J., and D.L. assisted with the experiments. E.M. and J.Z. assisted with the mouse model. F.Z., C.X., Z.W., C.X., J. Z., and J.L. provided insightful discussion and manuscript logic. G.W., Y.G., and J.L. analyzed data. G.W. and Y.G. prepared the manuscript with input from all authors.

## Data and material availability

All data generated in this study are included in the article and supplementary information files. Processed proteomics data are available in an Excel file named “Dataset S1_Analysis of Proteomics Data”. Processed bulk RNA-seq data are available in “Dataset S2_Analysis of Bulk-RNAseq Data”. Processed metabonomics data are available in “Dataset S3_Analysis of Metabonomics Data”. Processed scRNA-seq data are available in “Dataset S4_Analysis of scRNA-seq Data”. The RNAseq raw data and scRNAseq raw data have been deposited to the Gene Expression Omnibus (GEO) with accession numbers GSE280148 (token: ujurmeiabrivduz) and GSE280149 (token: ydabiwqwttodrab), respectively. Additional data, codes, and materials that support the findings of this research are available from the corresponding author upon reasonable request to the academic community.

## Supplementary Figure Legends

**Figure S1.**
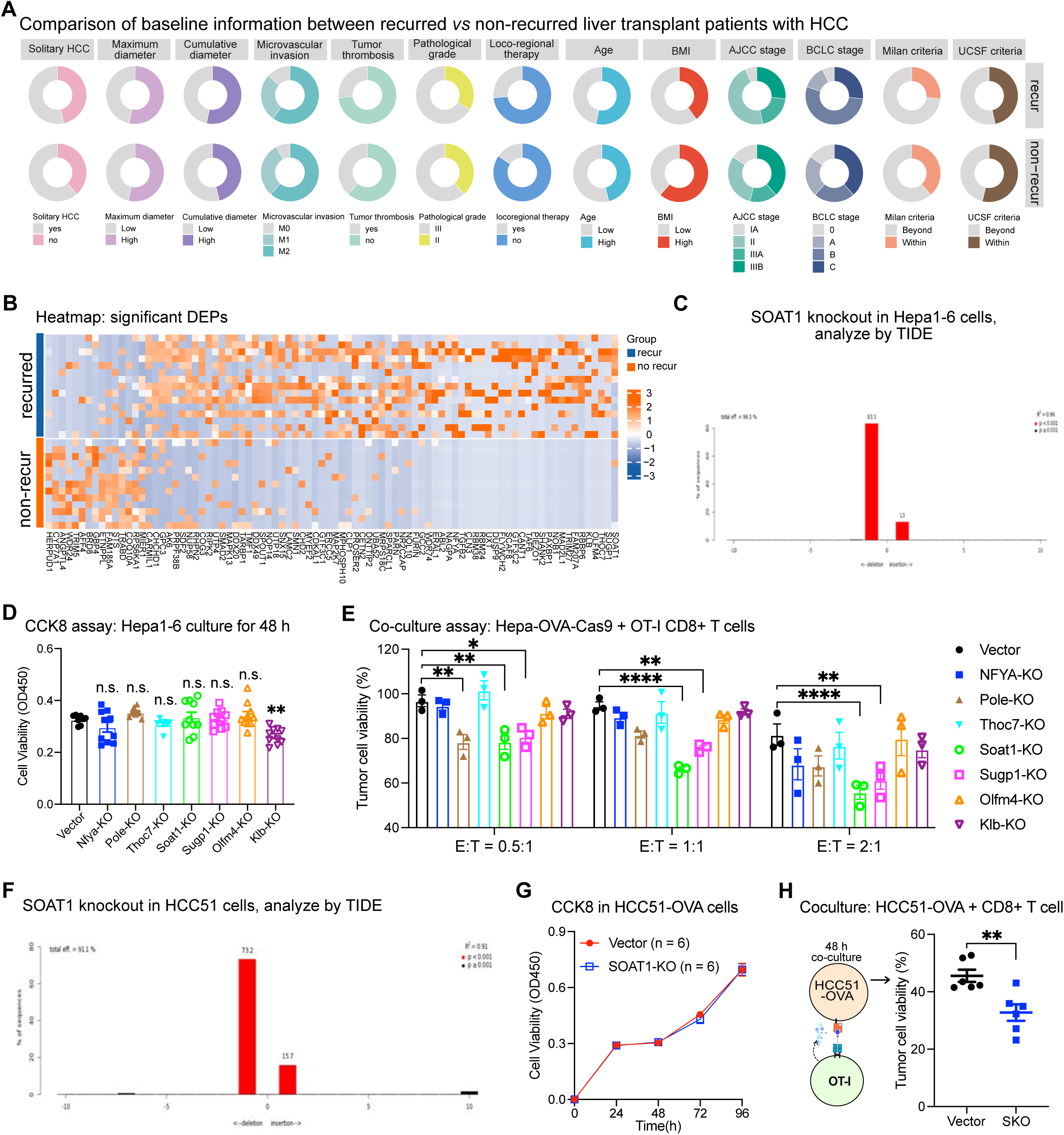
Differential proteomics-directed immune cell killing assays identifies SOAT1 as a key driver of HCC immune evasion and recurrence. (**A**) Baseline comparison of clinical characteristics between liver transplant patients with HCC who experienced recurrence versus those without later recurrence. (**B**) Heatmap of proteomics data depicting the differentially expressed proteins (DEPs, log2 FC ≥ 1.5 and P < 0.05) in the primary HCC tumors with later recurrence *versus* those without later recurrence. (**C**) SOAT1 knockout efficiency in Hepa1-6 cells was validated by targeted PCR and Sanger sequencing of the sgRNA-targeted genomic region. This was followed by TIDE (Tracking of Indels by Decomposition) analysis to assess indel formation. (**D**) *In vitro* proliferation of Hepa1-6 cells with individual knockout of target gene was assessed using the CCK-8 assay (n = 9). (**E**) Viability of Hepa-OVA-Cas9 cells with different gene knockouts after coculturing with OT-I CD8+ T cells at effector to target (E: T) ratios of 0.5: 1 and 2: 1 for 48 h (n =3). (**F**) SOAT1 knockout efficiency in HCC51 cells validated by Sanger sequencing of the targeted region, followed by TIDE analysis to confirm editing. (**G**) *In vitro* proliferation of SOAT1-KO and vector control HCC51 cells was assessed using the CCK-8 assay (n = 6). (**H**) Viability of SOAT1-KO and vector control HCC51-OVA cells after coculture with OT-I CD8+T cells at an E:T ratios of 2:1 for 48 h (n = 6). Data are shown as means ± SEMs; Statistical analysis was performed using a Student’s t test or two-way ANOVA. Asterisks: * P < 0.05, ** P < 0.01, *** P < 0.001, **** P < 0.0001. Related to Figure 1.

**Figure S2.**
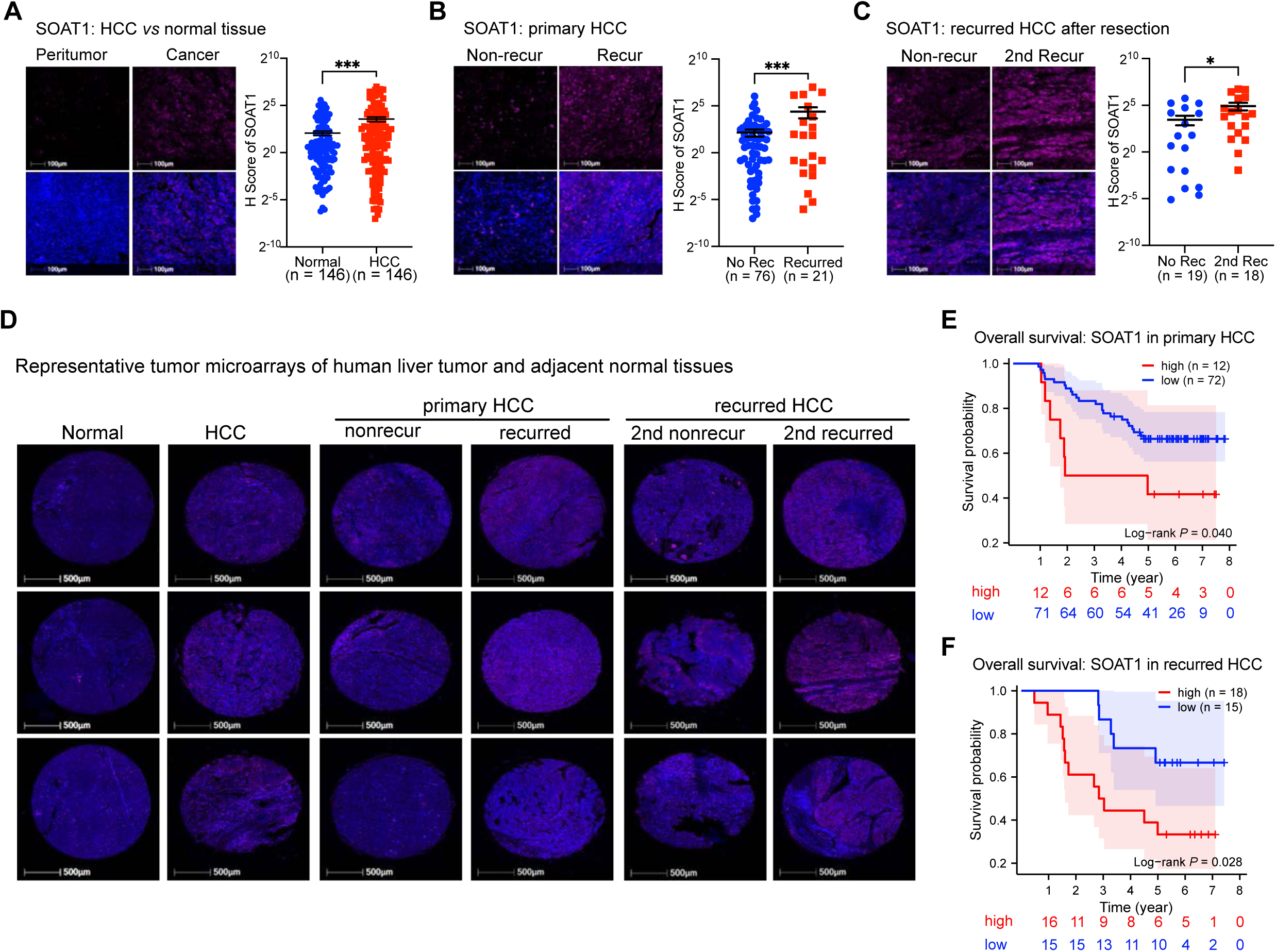
SOAT1 expression correlates with liver tumor recurrence and poor overall survival in liver transplant patients with HCC. (**A**) Representative immunofluorescence images (left) and quantification of SOAT1 expression (right) in paired human HCC tissues and adjacent normal tissues (n = 146). For human tissue microarrays (TMAs), SOAT1 expression was scored from 0 to 4 based on staining intensity using HALO software: negative, weak, moderate, or strong positive. The SOAT1 H-Score = [1×% SOAT1-weak cells + 2×% SOAT1-moderate cells + 3×% SOAT1-strong cells]. (**B**) Representative immunofluorescence images (left) and quantification of SOAT1 expression (right) in primary HCC tissues with recurrence (n = 21) compared to those without recurrence (n = 76) following liver transplantation. (**C**) Representative immunofluorescence images (left) and quantification of SOAT1 expression (right) in post-resection recurred HCC tissues with secondary recurrence (n = 18) after liver transplantation versus those without secondary recurrence (n = 19). (**D**) Representative immunofluorescence images from human tissue microarrays (TMAs) showing SOAT1 expression in paired human HCC tissues versus adjacent normal tissues, in primary HCC without recurrence and primary HCC with recurrence, as well as in recurred HCC without secondary recurrence and recurred HCC with secondary recurrence. (**E, F**) Clinical analysis of intra-tumoral SOAT1 expression and its impact on overall survival in patients with primary and post-resection recurred HCC. Patients with primary HCC, matched for disease stage, were divided into SOAT1-high (n = 12) and SOAT1-low (n = 72) cohorts using the average SOAT1 H-score as the cutoff. The SOAT1-high cohort exhibited significantly worse overall survival compared to the SOAT1-low cohort (**E**). Similarly, in patients with post-resection recurred HCC, intra-tumoral SOAT1-high cohort (n = 18) was also associated with significantly worse outcomes compared to the SOAT1-low cohort (n = 15) (**F**). Data are shown as means ± SEMs; Statistical analysis was performed using a Student’s t test or two-way ANOVA. Asterisks: * P < 0.05, ** P < 0.01, *** P < 0.001, **** P < 0.0001. Related to Figure 1

**Figure S3.**
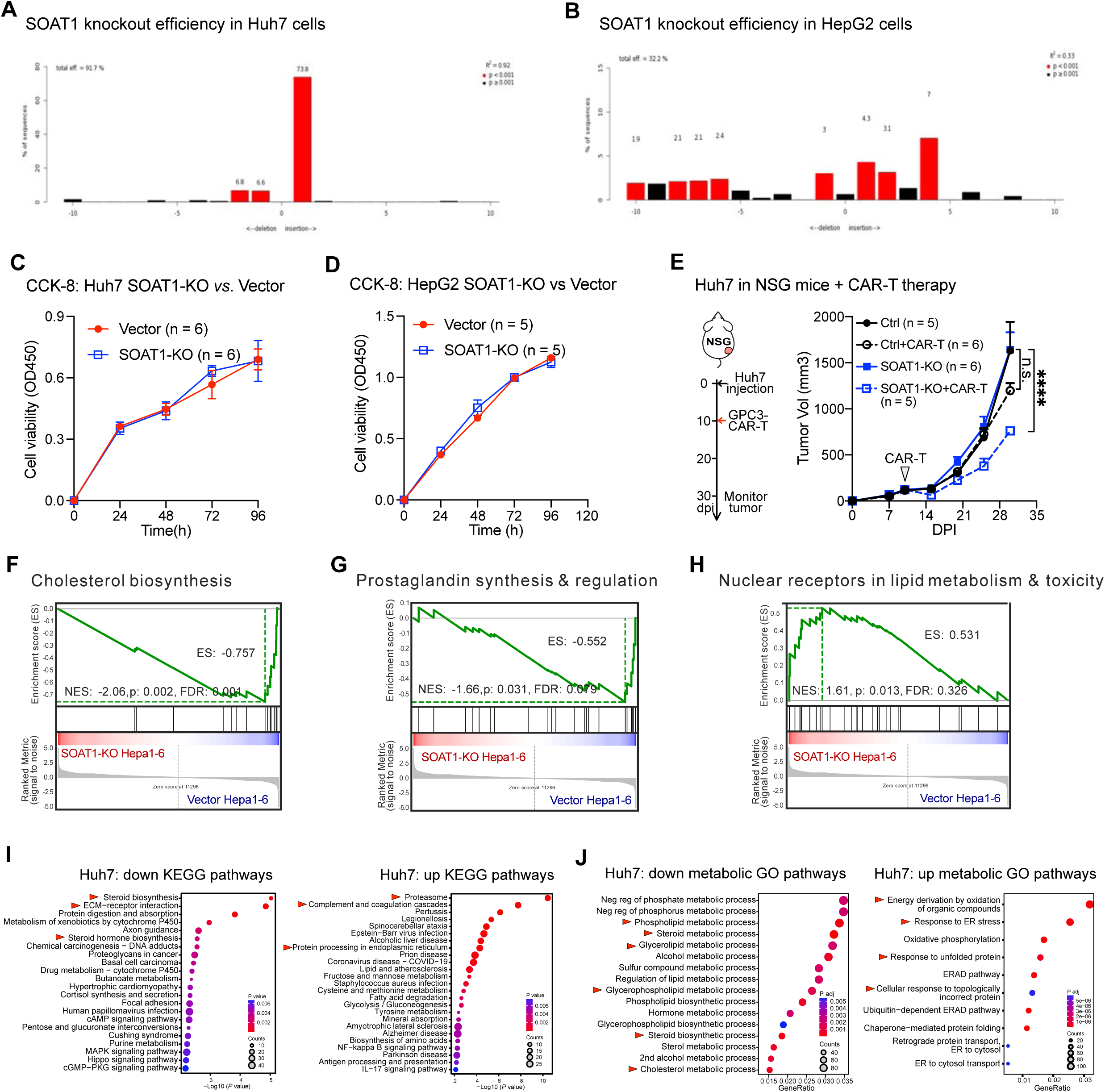
SOAT1 knockout sensitizes liver cancer cells to T cell-based immunotherapies. (**A, B**) SOAT1 knockout efficiency in Huh7 cells (**A**) and HepG2 cells (**B**) were confirmed by sanger sequencing of the sgRNA-targeted regions, followed by TIDE analysis. (**C, D**) *In vitro* proliferation of SOAT1-KO and vector control Huh7 cells (n = 6) (**C**), and SOAT1-KO and vector control HepG2 cells (n = 6) (**D**) were evaluated using the CCK-8 assay. (**E**) Schematics of CAR-T therapy *in vivo* (left panel). Growth curves of SOAT1-KO and vector control Huh7 tumors in immunodeficient NSG mice, treated with PBS or GPC3 CAR-T cells on day 10 post-tumor inoculation (n = 5 or 6 per group as indicated). (**F-H**) Gene Set Enrichment Analysis (GSEA) of the transcriptomes from SOAT1-KO versus vector Hepa1-6 cells shows significant downregulation of pathways related to cholesterol biosynthesis (**F**), prostaglandin synthesis and regulation (**G**), but upregulation of nuclear receptors in lipid metabolism and cytotoxicity (**H**). (**I**) KEGG pathway analysis reveals downregulated (left) and upregulated (right) pathways in SOAT1-KO Huh7 cells compared to vector controls. (**J**) Metabolic pathway analysis indicates downregulated (left) and upregulated (right) metabolic GO pathways in SOAT1-KO Huh7 cells versus vector controls. Data are shown as means ± SD; Statistical analysis was performed using a Student’s t test or two-way ANOVA. Asterisks: * P < 0.05, ** P < 0.01, *** P < 0.001, **** P < 0.0001. Related to Figure 2.

**Figure S4.**
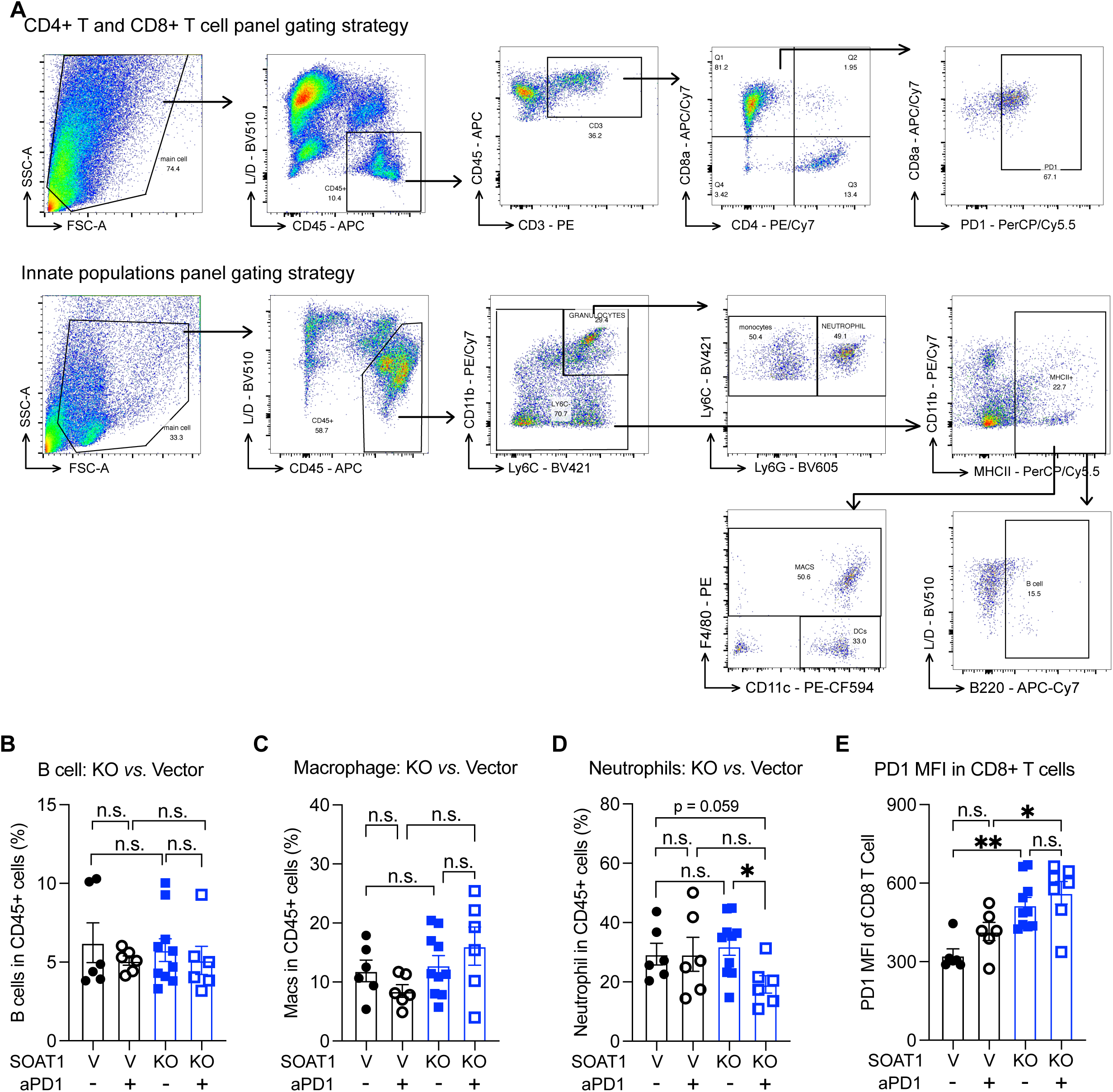
Flow cytometry to characterize tumor immune microenvironment fostered by SOAT1-KO and vector control Hepa1-6 tumors. (**A**) Gating strategies used in the flow cytometry analysis of adaptive immune cell populations (top) and innate immune populations (bottom) in tumor microenvironment. (**B-E**) Flow cytometry analysis of tumor immune microenvironment: quantification of B cells (**B**), macrophages (**C**), and neutrophils (**D**) among tumor-infiltrating CD45+ immune cells, and mean fluorescence intensity (MFI) of PD-1 expression in CD8+ T cells (**E**) in SOAT1-KO (n = 7 for anti-PD1, n = 10 for PBS) or vector control (n = 6) Hepa1-6 tumors in mice treated with PBS or anti-PD1at 30 days post transplantation. Data are shown as means ± SEMs; Statistical analysis was performed using a Student’s t test or two-way ANOVA. Asterisks: * P < 0.05, ** P < 0.01, *** P < 0.001, **** P < 0.0001. Related to Figure 3.

**Figure S5.**
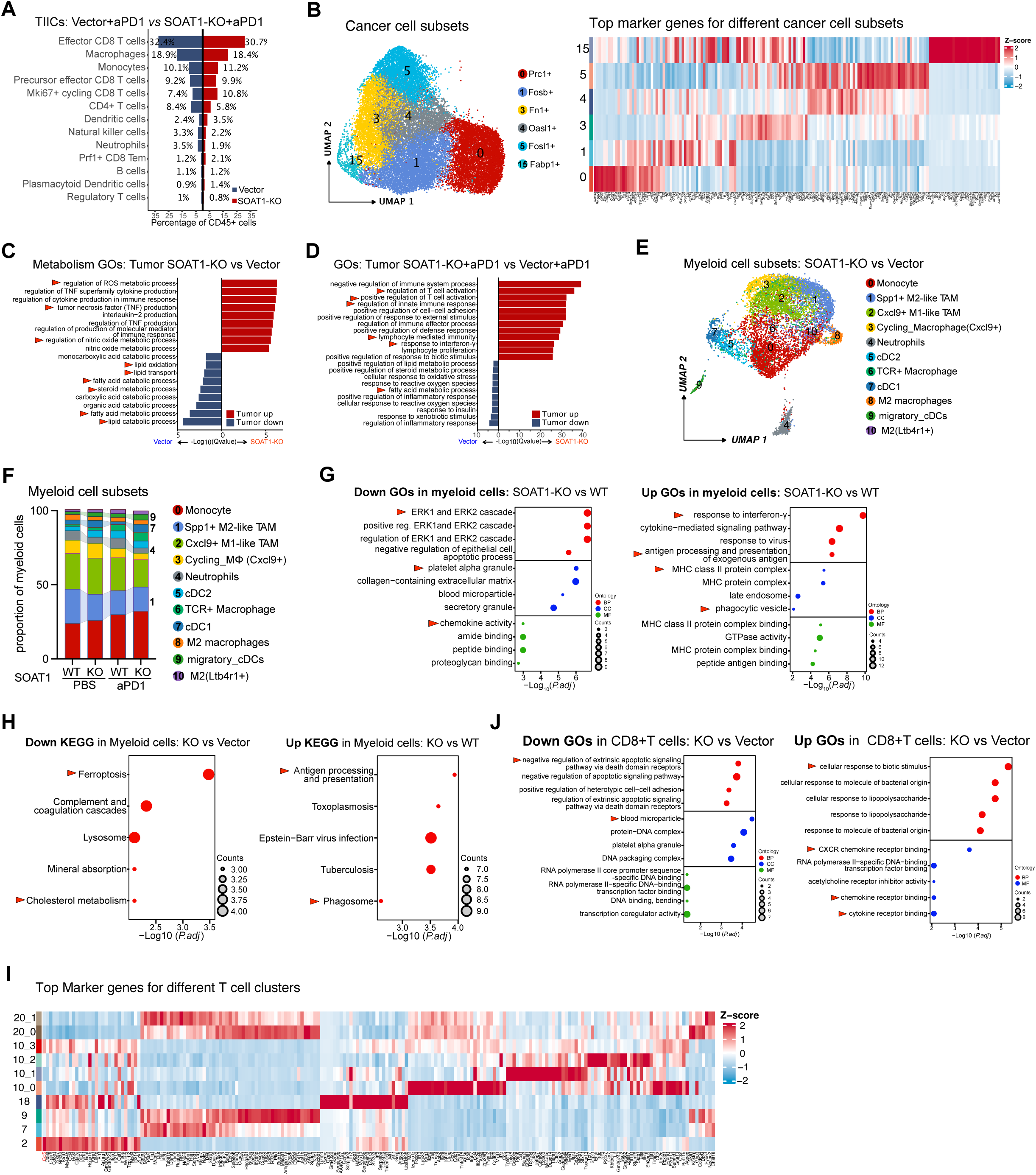
Single cell RNA-seq analysis of immune cell composition and transcriptomic profiles in SOAT1-KO and vector Hepa1-6 tumors. (**A**) Bar plots showing proportional differences in immune cell subsets among tumor-infiltrating CD45+ immune cells in SOAT1-KO tumors compared to vector control tumors in C57BL/6 mice treated with anti-PD1. (**B**) Top differentially expressed genes across 6 different clusters of Hepa1-6 cancer cells *in vivo*. (**C**) Metabolic GO analysis revealed the upregulated and downregulated pathways in SOAT1-KO versus vector Hepa1-6 cancer cells *in vivo,* based on scRNA-seq data. (**D**) GO analysis revealed the upregulated and downregulated pathways in SOAT1-KO versus vector Hepa1-6 cancer cells from anti-PD1 treatment group, based on scRNA-seq data. (**E**) UMAP plot of scRNA-seq profiles from SOAT1-KO and vector control Hepa1-6 tumors showing 11 myeloid cell clusters. (**F**) Barplot displaying proportional differences in myeloid cell subsets from SOAT1-KO versus vector control tumors in mice treated with PBS or anti-PD1 of scRNA-seq. (**G**) GO analysis of downregulated (left) and upregulated (right) pathways in tumor-infiltrated myeloid cells from SOAT1-KO versus vector control tumors in PBS-treated group. (**H**) KEGG analysis demonstrated the downregulated and upregulated pathways in tumor-infiltrated myeloid cells from SOAT1-KO versus vector control liver tumors from PBS treatment group. (**I**) Top differentially expressed genes among 10 different distinct clusters of T cells. (**J**) GO analysis revealed the downregulated (left) and upregulated (right) pathways in tumor-infiltrated CD8+T cells from SOAT1-KO tumors versus vector control tumors in the PBS groups. Data are shown as means ± SEMs; Statistical analysis was performed using a Student’s t test or two-way ANOVA. Asterisks: * P < 0.05, ** P < 0.01, *** P < 0.001, **** P < 0.0001. Related to Figure 3.

**Figure S6.**
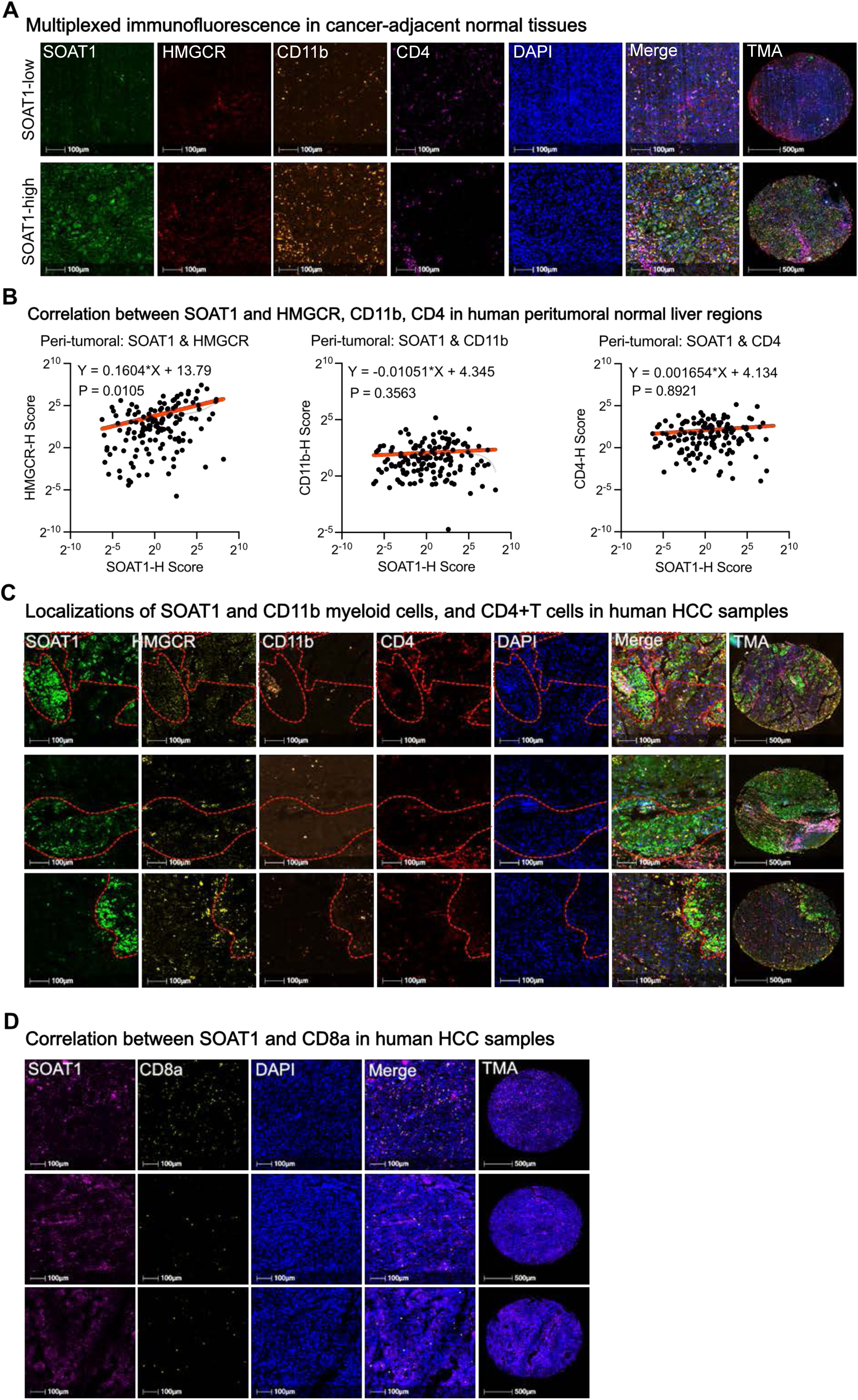
Multiplexed immunofluorescence staining demonstrates the relationship between SOAT1 expression, cholesterol metabolism, and myeloid cell infiltration in clinical HCC samples. (**A**) Representative multiplexed immunofluorescence images of TMAs showing the colocalization of SOAT1, HMGCR, CD11b, CD4, and DAPI in cancer-adjacent normal liver tissues. These images illustrate the spatial relationships between SOAT1 expression, cholesterol biosynthesis (HMGCR), CD11b+ myeloid cells, and CD4+ T cells, showcasing examples from HCC samples with low and high SOAT1 expression. (**B**) Correlation plots demonstrate the relationship between SOAT1 expression and HMGCR (left), CD11b (middle), or CD4 (right) levels in human peritumoral normal liver tissues. (**C**) Multiplexed immunofluorescence images of human HCC TMAs showing the distribution of SOAT1, HMGCR, CD11b+ myeloid cells, and CD4+ T cells in HCC tumors. (**D**) Multiplexed immunofluorescence images showing the localizations and relationships of SOAT1 and CD8+ T cells in human HCC tissues. Data are shown as means ± SEMs; Statistical analysis was performed using a Student’s t test or two-way ANOVA. Asterisks: * P < 0.05, ** P < 0.01, *** P < 0.001, **** P < 0.0001. Related to Figure 3.

**Figure S7.**
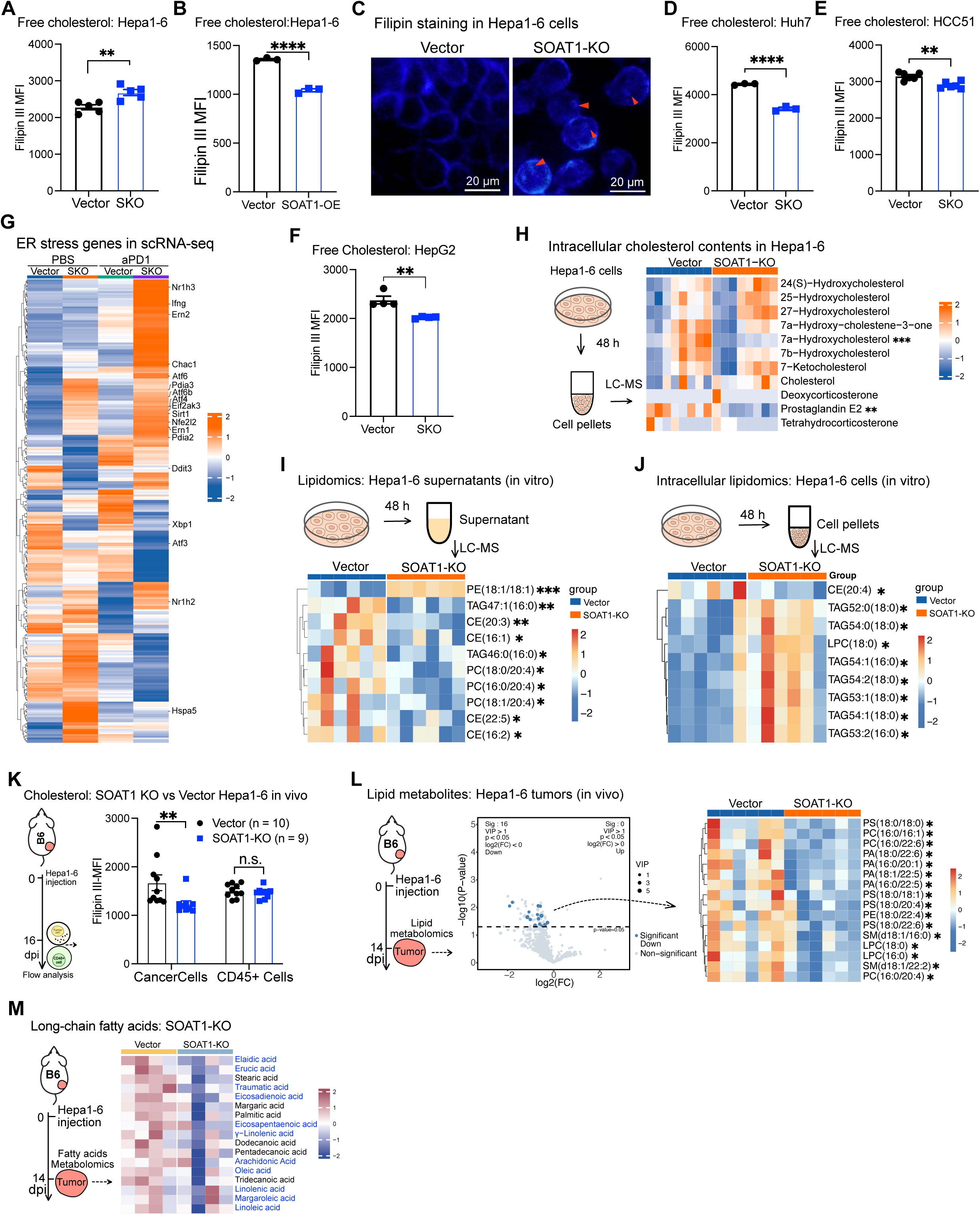
SOAT1 knockout inhibits cholesterol and fatty acid biosynthesis and metabolism. (**A**) Cholesterol distribution in SOAT1-KO and vector control Hepa1-6 cells: Total cellular total free cholesterol levels were quantified using Filipin III staining (n = 5). (**B**) Filipin III staining shows changes in subcellular cholesterol localization after SOAT1 knockout in Hepa1-6 cells. (**C**) Total cellular free cholesterol levels in SOAT1-KO and vector control Huh7 cells were quantified using Filipin III staining. (**D**) Total cellular free cholesterol levels were assessed via Filipin III staining in SOAT1-overexpressing (SOAT1-OE) and vector control Hepa1-6 cells (n = 3). (**E-F**) Cholesterol distribution in SOAT1-KO and vector control cells: Total cellular total free cholesterol levels were quantified using Filipin III staining in HCC51(**E**) (n = 4) and HepG2 (**F**) (n = 6) cells. (**G**) Heatmap illustrating the differential expression of ER stress-related genes in SOAT1-KO versus vector control Hepa1-6 tumor cells from C57BL/6 mice treated with PBS or anti-PD1, according to scRNA-seq data. (**H**) Heatmap displays intracellular cholesterol metabolites in the *in vitro* cultured SOAT1-KO and vector Hepa1-6 cells (n = 8). (**I-J**) Lipid-targeted metabolomics: heatmap showing differential lipid metabolites in the supernatant (**I**) and cell pellets (**J**) of SOAT1-KO versus vector Hepa1-6 cells cultured *in vitro* (n = 6). These data indicated decreased cholesteryl esters (CE) and phosphocholine (PC), while triacylglycerols (TAG) were increased in SOAT1-KO cells. (**K**) Total cellular free cholesterol levels in SOAT1-KO and vector Hepa1-6 cancer cells, as well as in tumor-infiltrating CD45+ immune cells (TIICs), were analyzed by Filipin III staining. Cancer cells and TIICs were isolated from C57BL/6 mice 14 days post-inoculation (n = 10). (**L**) Lipid-targeted metabolomics *in vivo*: volcano plot (left) and heatmap (right) showing differential lipid metabolites in SOAT1-KO and vector control Hepa1-6 tumors from mice 14 days post-inoculation (n = 6). (**M**) Heatmap depicting differences in intracellular long-chain fatty acids between SOAT1-KO and vector control Hepa1-6 tumors, with unsaturated fatty acids labelled in blue (n = 4). Data are shown as means ± SEMs; Statistical analysis was performed using a Student’s t test or two-way ANOVA. Asterisks: * P < 0.05, ** P < 0.01, *** P < 0.001, **** P < 0.0001. Related to Figure 4.

**Figure S8.**
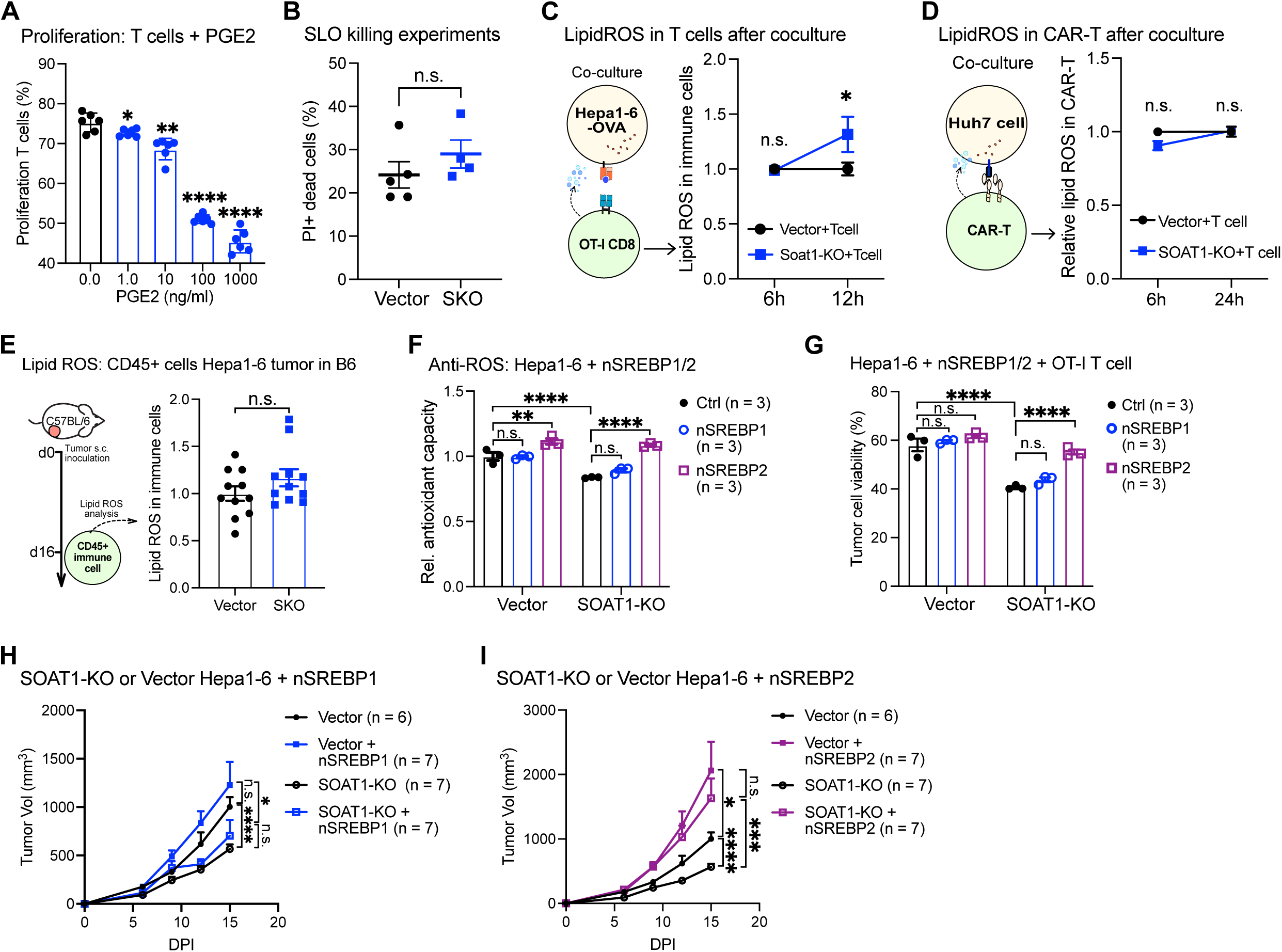
SOAT1 knockout sensitizes liver cancer cells to T cell-mediated cytotoxicity by reducing cellular antioxidant capacity. (**A**) The effect of PGE2 on *ex vivo* T cell proliferation was assessed by adding different concentrations of PGE2 into CFSE-labelled CD8+T cells primed with anti-CD3/anti-CD28 (n = 6). (**B**) Streptolysin O (SLO) permeabilization assay: plasma membrane resealing after SLO permeabilization was assessed using propidium iodide (PI) staining. Flow cytometry analysis showed no significant difference in PI+ dead cells between SOAT1-KO (n = 5) and vector (n = 4) Hepa1-6 cells. (**C, D**) Lipid ROS levels in OT-I CD8+ T cells after coculture with SOAT1-KO and vector Hepa-OVA-Cas9 cells for 6 and 12 h (**C**), and in GPC3 CAR-T cells cocultured with SOAT1-KO and vector Huh7 cells for 6 and 24 h (**D**), were evaluated using lipid peroxidation sensor BODIPY-C11 (n = 5 in 6h sampling, n = 4 in 12h sampling). (**E**) Lipid ROS levels in tumor-infiltrated CD45+ immune cells from SOAT1-KO and vector Hepa1-6 tumors in immunocompetent C57BL/6 mice, assessed by BODIPY-C11 staining (n = 11). (**F**) The anti-oxidant capacity of control, nSREBP1, or nSREBP2-overexpressed SOAT1-KO and vector Hepa-OVA-Cas9 cells (n = 3). (**G**) Viability of control, nSREBP1, or nSREBP2-overexpressed SOAT1-KO and vector Hepa-OVA-Cas9 cells when cocultured with OT-I CD8+ T cells for 48 h (n = 3). (**H, I**) Growth curves of subcutaneously transplanted SOAT1-KO and vector Hepa1-6 tumors in C57BL/6 mice, after re-expressing the transcriptional active form of N-terminal SREBP1 (nSREBP1) (**H**), active form of N-terminal SREBP2 (nSREBP2) (**I**). Data are shown as means ± SEMs; Statistical analysis was performed using a Student’s t test or two-way ANOVA. Asterisks: * P < 0.05, ** P < 0.01, *** P < 0.001, **** P < 0.0001. Related to Figure 5.

**Figure S9.**
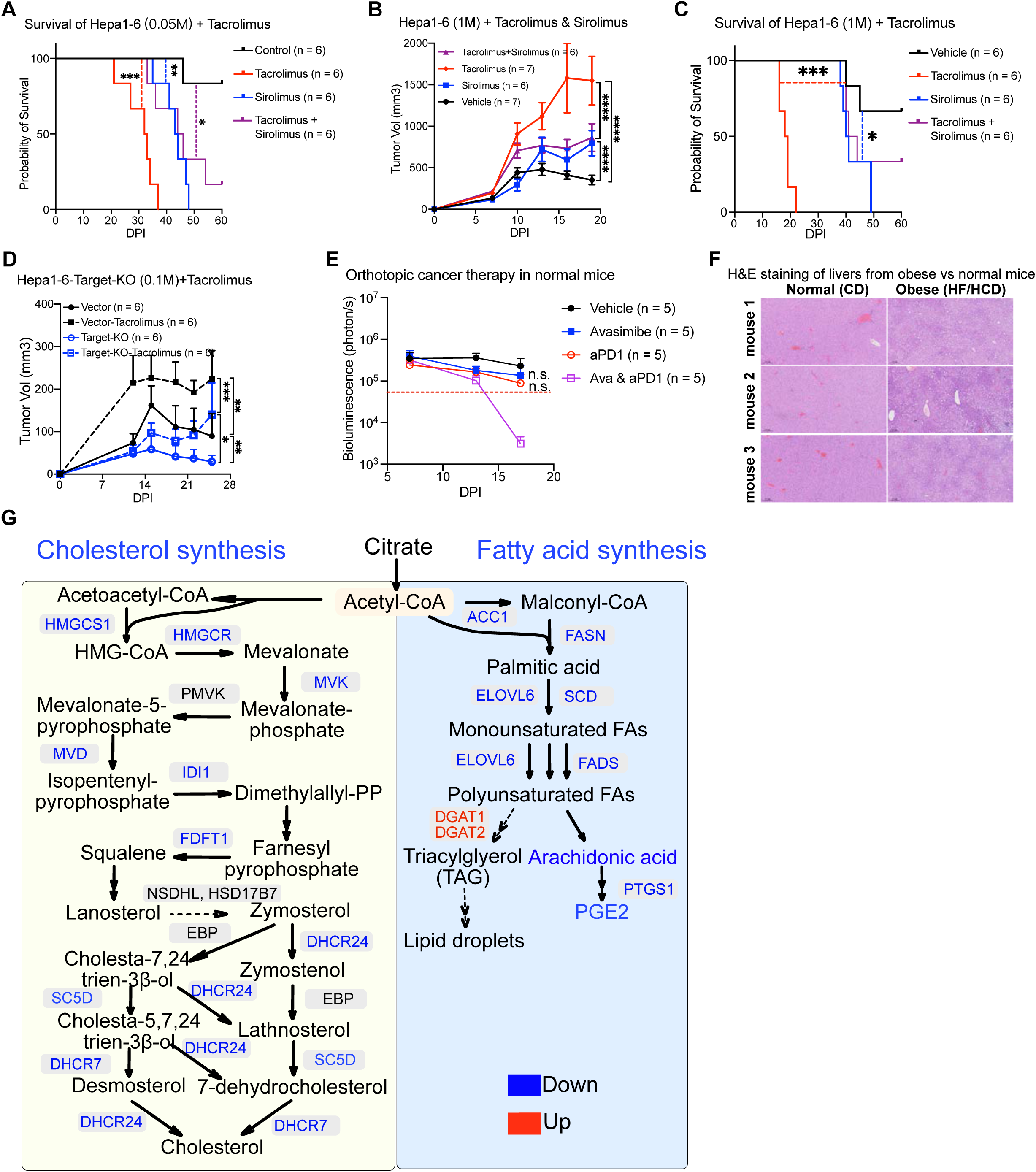
SOAT1 knockout suppresses liver cancer growth and enhances the efficacy of CAR-T therapy in conditions of suboptimal immunosurveillance. (**A**) Kaplan-Meier survival curves of C57BL/6 mice subcutaneously inoculated with 5×10^4^ Hepa1-6 cells and treated with PBS, sirolimus, tacrolimus, and their combination (n = 6). (**B, C**) Tumor growth curves (**B**) and Kaplan-Meier survival curves (**C**) of C57BL/6 mice subcutaneously inoculated with 5×10^4^ Hepa1-6 cells, treated with PBS, sirolimus, tacrolimus, or a sequential combination of tacrolimus followed by sirolimus (n = 6). (**D**) Growth curves of subcutaneous Hepa1-6 tumor in C57BL/6 mice inoculated with 1×10^5^ SOAT1-KO or vector control Hepa1-6 cells, treated with either PBS or tacrolimus (n = 6). (**E**) Growth of orthotopic liver tumors in common diet (CD)-fed mice, assessed by measuring luciferase activity. Mice were treated with vehicle, avasimbe, anti-PD1, or their combination starting on day 10 post-tumor inoculation (n = 5). (**F**) Representative hematoxylin and eosin (H&E) staining images comparing liver tissues from obese mice, fed with a high-fat, high-cholesterol diet (HF/HCD), versus normal diet-fed mice. Liver tissues from obese mice exhibit increased steatosis and lipid accumulation, contrasting with normal liver histology. (**G**) Schematics illustrating the enzymes downregulated (blue) or upregulated (red) in cholesterol and fatty acid biosynthesis following SOAT1 knockout in Hepa1-6 and Huh7 cells, based on the bulk and single-cell RNA-seq data. Data are shown as means ± SEMs; Statistical analysis was performed using a Student’s t test or two-way ANOVA. Asterisks: * P < 0.05, ** P < 0.01, *** P < 0.001, **** P < 0.0001. Related to Figure 6.

## Supplementary Materials

Materials and Methods

Figs. S1 to S9

Tables S1 and S2

Dataset S1 to S4

## Supplemental Tables

Table S1. Baseline characteristics of liver transplant patients with HCC: comparison between recurrence and non-recurrence groups.

Table S2. Baseline characteristics of HCC patients: comparison between obese group and normal body weight group.

## Supplemental Datasets

Dataset S1. Analysis of Proteomics

Data Dataset S2. Analysis of Bulk-RNAseq

Data Dataset S3. Analysis of Metabonomics

Data Dataset S4. Analysis of scRNAseq Data

## References

1. Llovet, J. M. et al. Hepatocellular carcinoma. Nat. Rev. Dis. Primer 7, 1–28 (2021).

2. Marsh, J. W. et al. The prediction of risk of recurrence and time to recurrence of hepatocellular carcinoma after orthotopic liver transplantation: A pilot study. Hepatology 26, 444 (1997).

3. Zimmerman, M. A. et al. Recurrence of Hepatocellular Carcinoma Following Liver Transplantation: A Review of Preoperative and Postoperative Prognostic Indicators. Arch. Surg. 143, 182–188 (2008).

4. Forner, A., Llovet, J. M. & Bruix, J. Hepatocellular carcinoma. The Lancet 379, 1245–1255 (2012).

5. Sharma, P., Hu-Lieskovan, S., Wargo, J. A. & Ribas, A. Primary, Adaptive, and Acquired Resistance to Cancer Immunotherapy. Cell 168, 707–723 (2017).

6. Hegde, P. S. & Chen, D. S. Top 10 Challenges in Cancer Immunotherapy. Immunity 52, 17–35 (2020).

7. Ringel, A. E. et al. Obesity Shapes Metabolism in the Tumor Microenvironment to Suppress Anti-Tumor Immunity. Cell 183, 1848–1866.e26 (2020).

8. Tsai, C.-H. et al. Immunoediting instructs tumor metabolic reprogramming to support immune evasion. Cell Metab. 35, 118–133.e7 (2023).

9. Faubert, B., Solmonson, A. & DeBerardinis, R. J. Metabolic reprogramming and cancer progression. Science 368, eaaw5473 (2020).

10. Rathmell, J. C. Obesity, Immunity, and Cancer. N. Engl. J. Med. 384, 1160–1162 (2021).

11. Sung, H. et al. Global patterns in excess body weight and the associated cancer burden. CA. Cancer J. Clin. 69, 88–112 (2019).

12. Body-mass index and incidence of cancer: a systematic review and meta-analysis of prospective observational studies - The Lancet. https://www.thelancet.com/journals/lancet/article/PIIS0140-6736(08)60269-X/fulltext.

13. Hoy, A. J., Nagarajan, S. R. & Butler, L. M. Tumour fatty acid metabolism in the context of therapy resistance and obesity. Nat. Rev. Cancer 21, 753–766 (2021).

14. Younossi, Z. et al. Nonalcoholic Steatohepatitis Is the Fastest Growing Cause of Hepatocellular Carcinoma in Liver Transplant Candidates. Clin. Gastroenterol. Hepatol. 17, 748–755.e3 (2019).

15. Estes, C., Razavi, H., Loomba, R., Younossi, Z. & Sanyal, A. J. Modeling the epidemic of nonalcoholic fatty liver disease demonstrates an exponential increase in burden of disease. Hepatology 67, 123 (2018).

16. Marengo, A., Rosso, C. & Bugianesi, E. Liver Cancer: Connections with Obesity, Fatty Liver, and Cirrhosis. Annu. Rev. Med. 67, 103–117 (2016).

17. Pardoll, D. M. The blockade of immune checkpoints in cancer immunotherapy. Nat. Rev. Cancer 12, 252–264 (2012).

18. Ribas, A. & Wolchok, J. D. Cancer immunotherapy using checkpoint blockade. Science 359, 1350–1355 (2018).

19. June, C. H., O’Connor, R. S., Kawalekar, O. U., Ghassemi, S. & Milone, M. C. CAR T cell immunotherapy for human cancer. Science 359, 1361–1365 (2018).

20. Llovet, J. M. et al. Immunotherapies for hepatocellular carcinoma. Nat. Rev. Clin. Oncol. 19, 151–172 (2022).

21. Kruse, B. et al. CD4+ T cell-induced inflammatory cell death controls immune-evasive tumours. Nature 1–8 (2023) doi:10.1038/s41586-023-06199-x.

22. Chang, T. Y., Chang, C. C. Y. & Cheng, D. ACYL-COENZYME A:CHOLESTEROL ACYLTRANSFERASE. Annu. Rev. Biochem. 66, 613–638 (1997).

23. Guan, C. et al. Structural insights into the inhibition mechanism of human sterol O-acyltransferase 1 by a competitive inhibitor. Nat. Commun. 11, 2478 (2020).

24. Morotti, M. et al. PGE2 inhibits TIL expansion by disrupting IL-2 signalling and mitochondrial function. Nature 1–9 (2024) doi:10.1038/s41586-024-07352-w.

25. Lacher, S. B. et al. PGE2 limits effector expansion of tumour-infiltrating stem-like CD8+ T cells. Nature 1–9 (2024) doi:10.1038/s41586-024-07254-x.

26. Liu, Z. et al. CPT1A-mediated fatty acid oxidation confers cancer cell resistance to immune-mediated cytolytic killing. Proc. Natl. Acad. Sci. 120, e2302878120 (2023).

27. De Martino, M., Rathmell, J. C., Galluzzi, L. & Vanpouille-Box, C. Cancer cell metabolism and antitumour immunity. Nat. Rev. Immunol. 1–16 (2024) doi:10.1038/s41577-024-01026-4.

28. Brewin, J. et al. Generation of EBV-specific cytotoxic T cells that are resistant to calcineurin inhibitors for the treatment of posttransplantation lymphoproliferative disease. Blood 114, 4792–4803 (2009).

29. Huang, Y. et al. A pan-cancer analysis identifies SOAT1 as an immunological and prognostic biomarker. Oncol. Res. 31, 193–205 (2023).

30. Jiang, Y. et al. Proteomics identifies new therapeutic targets of early-stage hepatocellular carcinoma. Nature 567, 257–261 (2019).

31. Li, Y. et al. 7-Dehydrocholesterol dictates ferroptosis sensitivity. Nature 1–8 (2024) doi:10.1038/s41586-023-06983-9.

32. Freitas, F. P. et al. 7-Dehydrocholesterol is an endogenous suppressor of ferroptosis. Nature 1–10 (2024) doi:10.1038/s41586-023-06878-9.

33. Meiner, V. L. et al. Disruption of the acyl-CoA:cholesterol acyltransferase gene in mice: Evidence suggesting multiple cholesterol esterification enzymes in_mammals. Proc. Natl. Acad. Sci. 93, 14041–14046 (1996).

34. Yilmaz, S. & Ince, V. The Importance of the Immunosuppressive Regime on Hepatocellular Carcinoma Recurrence After Liver Transplantation. J. Gastrointest. Cancer 52, 1350–1355 (2021).

35. Yang, W. et al. Potentiating the antitumour response of CD8 + T cells by modulating cholesterol metabolism. Nature 531, 651–655 (2016).

