## Supplementary material for "Cholesterol esterification blockade boosts immunotherapy and reduces cancer relapse": methods

### Materials and Methods

**Cell lines.**

HEK293FT cell line (Thermo, Cat# R70007) was purchased from Thermo. Mouse hepatoma cell line Hepa1-6 were kindly provided by Dr. Chenqi Xu Lab at CEMCS. Mouse hepatocellular carcinoma cell line HCC51 was established in Guangchuan Wang Lab at CEMCS by isolating and culturing the primary hepatocellular carcinoma in AAV-TBG-Cre injected CAG-LSL-MYC, P53fl/fl, ROSA26-LSL-Cas9 mice on a C57BL/6 background. Human hepatoma cell line Huh-7 and human hepatocellular carcinoma cell line HepG2 was purchased from cell bank of Chinese Academy of Sciences Center for Excellence in Molecular Cell Science (CEMCS, CAS). All the purchased cell lines have been authenticated by the original vendors. These cell lines were cultured in Dulbecco's minimal essential medium (DMEM) supplemented with 8% fetal bovine serum (FBS) and 1% penicillin/streptomycin, in a $CO_2$ cell incubator 37°C.

**Mice.**

All mice experiments were conducted in compliance with the ethical guidelines of the Institutional Animal Care and Use Committee (IACUC) of Chinese Academy of Sciences Center for Excellence in Molecular Cell Science (CEMCS, CAS), or Shanghai medical college, Fudan University. C57BL/6 inbred mice were purchased from JiHui or LingChang (Shanghai, China), and B-NDG mice were purchased from Biocytogen (Shanghai, China). $Rag2^{-/-}$ transgenic mice were kindly gifted by Dr. Xiaolong Liu (CEMCS, CAS) and OT-I TCR transgenic mice were kindly gifted by Dr. Hongyan Wang (CEMCS, CAS). Mice at the ages of 6 to 9 weeks were used for experiments, and were housed in specific pathogen-free animal facilities at CEMCS, CAS or Shanghai medical college. Mice were randomized to receive different treatments based on age, weight, and tumor sizes, typically 5-10 mice per group.

**Proteomics analysis of clinical liver cancer samples**

All human samples involved in this study were performed in accordance with the principles outlined in the 2013 revised Declaration of Helsinki and was approved by the Ethics Committee of Huashan Hospital, Fudan University (KY-2019–511 and KY2024-595). Written informed consent was obtained from all participants. The proteomic sequencing of 28 HCC specimens obtained from the Liver Transplantation Center of Huashan Hospital, Fudan University, was performed by the National Institute of Metrology (China) as described below. Proteomic analysis of human samples was performed in four key steps: protein isolation from formalin-fixed paraffin-embedded (FFPE) tumor samples, protein digestion, peptide sequencing via mass spectrometry (MS), and data analysis to identify proteins.

Detailed processing of the MS raw files as follows: the MS raw files were searched in the NCBI Human Refseq database using the Mascot search engine (v2.3, Matrix Science Inc.; database version 04-07-2013, containing 32015 entries). The mass deviations for parent and daughter ions were set at 20 and 50 ppm, respectively (Q Exactive HF) or 0.5 Da. Protein cleavage sites were defined as arginine (R) and lysine (K), allowing up to two missed cleavages. Carbamidomethylation (C) for immobilization and acetyl (protein N-term) and oxidation (M) for dynamic modifications. All the identified peptides were obtained from the area under the MS1 peak calculation, with a false discovery rate controlled at 1%. At the protein characterization level, only those proteins with at least one unique peptide and two high-quality peptides (strict peptide, i.e., Mascot Ion Score >20) were retained. For protein quantification, we used an intensity-based absolute quantification algorithm (i.e., iBAQ algorithm) and normalized each iBAQ value to the Fraction of Total (FOT-iBAQ) by dividing the iBAQ value by the sum of the iBAQ values for all detected proteins in the corresponding sample. To make this FOT size easier to read and write, the FOT values were multiplied by $10^5$, resulting in the iFOT value.

**Clinical data collection and analysis**

From February 2016 to November 2018, a total of 153 patients with hepatocellular carcinoma (HCC) who underwent Liver transplantation (LT) at Huashan Hospital, Fudan University, Shanghai, China, were included in the study. All transplanted livers were matched through the China Organ Transplant Response System (COTRS) and sourced from cardiac death donors through Organ Procurement Organizations (OPO). All patients were followed up until June 2024 or until relapse occurred. Inclusion criteria for the study were HCC patients without extrahepatic metastasis and macrovascular invasion. Exclusion criteria included the presence of other malignant tumors, death within three months post-surgery, and incomplete follow-up data.

Patient data were collected by consulting patients' admission records, medical histories and follow-ups from Huashan hospital. During the first three months of follow-up after LT, patients were examined weekly. In the following three months, follow-ups were scheduled bi-weekly, and monthly thereafter for subsequent six months. During each follow-up, an ultrasound examination of the liver was conducted. For the first two years post-LT, computed tomography and liver enhancement magnetic resonance imaging were performed every three months. For patients with HCC recurrence, a multidisciplinary team provided regional therapies (e.g. surgery, radiofrequency ablation) or systemic treatments (e.g. targeted therapies, chemotherapy) along with supportive care. Recurrence-free survival (RFS) was defined as the time from surgery to either HCC occurrence including intrahepatic recurrence or distant metastasis, or the end of follow-up. According to the inclusion and exclusion criteria, actual follow-up status, and availability of complete data from pathological specimens, 131 patients were eligible for the overall survival (OS) analysis, and 118 patients were eligible for RFS analysis. Tumor characterizations, including number, size, pathological grade, and other histopathological features, were obtained from pathological examinations of the recipient liver specimens resected during LTs.

**Cholesterol and lipid targeted metabolomics by LC-MS analysis**

SOAT1-KO and vector control Hepa1-6 cells were cultured in DMEM supplemented with 8% FBS for 48 hours. After culture, the supernatant and cell pellets were collected and stored at -80°C before being used for subsequent analysis of cholesterol metabolites, lipid metabolites, and long-chain fatty acids using targeted UPLC-ESI-MS/MS or LC-MS/MS in Shanghai Oebiotech Co., Ltd. For direct examination of *in vivo* tumors, SOAT1-KO and vector control Hepa1-6 tumors were harvested from C57BL/6 mice 14 days after subcutaneously inoculation. The tumor samples were flash-frozen in liquid nitrogen and stored at -80°C. The preserved serum samples were subsequently sent to Oebiotech (Shanghai) for UPLC-ESI-MS/MS of cholesterol and its derivatives.

**Isolation and culture of mouse naïve CD8+ T cells**

Mouse naive $CD8^+$ T cells were isolated from OT-I mice using mouse naive $CD8^+$ T cell isolation kit (StemCell, Cata#19858). Spleens and lymph nodes of OT-I mice were collected, smashed and filtered using a 70 μm cell strainer. After twice washing with MACS buffer (0.5% BSA, 2 mM EDTA in PBS), red blood cells (RBCs) were lysed with RBC lysis buffer for 3-5 min at room temperature and then diluted with 10X volume of PBS before centrifugation at 300 g for 5 min at 4°C. The splenocytes were washed twice with MCAS buffer before naive $CD8^+$ T cells were isolated using naïve $CD8^+$ T isolation kits (Stem cell). Naive $CD8^+$ T cells activated by 5 μg/ml plate-coated anti-CD3 and 1 μg/ml soluble anti-CD28, otherwise the splenocytes were directly activated by adding SIINFEKL peptide for 2-3 days in R10 supplemented with 2 ng/ml IL-2, 5 ng/ml IL-7, and 5 ng/ml IL-15 at 37°C.

**Cellular cholesterol and lipid measurement.**

Filipin III staining, ALOD4-AF647 staining, and BODIPY493/503 staining were used to measure total cellular free cholesterol, accessible plasma membrane cholesterol, and cellular neutral lipid content, respectively. To measure the total free cholesterol, SOAT1-KO or vector control Hepa1-6 cells, HCC51 cells, Huh7 cells, HepG2 cells, or single-cell suspensions of tumor were stained with flow antibodies at 4°C for 30 min, washed and fixed with 4% paraformaldehyde for 10 min at room temperature. After washing off the PFA, cells were resuspended in 50 μg/ml Filipin III (Biogems, Cat# 4804999) for 30 min at room temperature. To assess accessible plasma membrane cholesterol, cells were incubated with 20 mg/ml ALOD4-AF647 in RPMI 1640 at 37°C for 30 min, before being stained with flow antibodies at 4°C for 30 min if applicable[2]. After staining, cells were washed and, if necessary, stained with flow antibodies at 4°C for an additional 30 min. All stained cells were washed twice or three times with MACS buffer before being analyzed by flow cytometry.

**Cell proliferation assay**

Cell viability and proliferation was assessed using a Cell Counting Kit-8 (CCK-8) assay kit (Vazyme, China). A total of 0.5-1 × $10^4$ cells per well were cultured in triplicate in a 96-well microplate (Corning, USA). After 24, 48, 72, and 96 h of treatments or culture, 10 μl of CCK-8 reagent (diluted in 100 μl of serum-free medium) was added to each well, followed by a 3-hour incubation at 37 °C. Absorbance at 450 nm was measured using a multifunction enzyme-linked analyzer (Thermo Fisher) to calculate cell viability.

**Lentivirus production.**

Lentiviruses were generated using 3-plamid system by co-transfecting HEK293FT cells (Thermo, Cat# R70007). Briefly, HEK293FT cells were seeded in 150-mm dishes one day prior to transfection, ensuring they reached 80-90% confluency at the time of transfection. Upon reaching the desired confluency, the medium was replaced with serum-free DMEM. A mixture of three plasmids and polyethyleneimine (PEI) were prepared by adding 20 μg of lentiviral expression plasmid, 15 μg of psPAX2, and 10 μg of pMD2.G into 130 μL OptiMEM, followed by thoroughly mixing. To this mixture, 130 μL of 1 mg/ml PEI was added, mixed well, and incubated at room temperature for 10-15 min. The mixture was then added dropwise into HEK293FT cells. Six hours after transfection, the medium was replaced with DMEM supplemented with 8% FBS. Lentiviruses-containing supernatants were collected at 48 h and 72 h post-transfection, centrifuged at 800 g for 5 min, and filtrated through a 0.45-μm filter to remove debris. The lentiviruses were then concentrated by ultracentrifugation at 27,000 rpm for 2h at 4°C, resuspended in serum-free medium, and stored at −80 °C until use.

**CAR-T cell production**

Peripheral blood mononuclear cells (PBMCs) were isolated from heathy donors using the Ficoll-Paque density gradient centrifugation method. Briefly, Ficoll-Paque Plus was warmed to room temperate and aliquoted into 15 ml conical tubes, with 5 ml per tube. 10 ml of PBS diluted whole blood (1:1) was carefully overlayered on top of the Ficoll-Plaque Plus and centrifuge at 400 g for 35 min at room temperature without using brakes. The buffy layer containing PBMCs at Ficoll-plasma interface were carefully collected, diluted with five volume of PBS and centrifuged again. The PBMCs were washed, counted, aliquoted, and frozen at a density of 1-5×$10^7$ cells per vial using freezing medium (10% DMSO and 90% FBS).

For CAR-T production, T cells within PBMCs were either sorted by FACS or directly activated with 12.5 µl/ml ImmunoCult™ Human CD3/CD28 T Cell Activator (StemCell, Cat#10971) for 3 days in human T cell medium. The human T cell medium consisted of X-VIVO medium (Lonza, Cat#04-418Q) supplemented with 5% FBS, 2 mM L-glutamine, 49 nM 2-mercaptoethanol, 10 mM N-acetyl-l-cysteine, 2 ng/mL hIL-2, 5 ng/ml hIL-7 and 5 ng/ml hIL-15. After 2-5 days of activation, T cells were collected, washed and seeded at a density of $1\times10^6$ cells per well in 24-well plate. Polybrene (0.8 µg/ml) and concentrated lentiviruses were then added, and the cells were gently mixed and spun at 900 g for 90 min at 32°C. Following spin-infection, T cells were cultured at 37°C for 16-24 h before replacing the virus-containing medium was fresh human T cell medium. CAR-T cells were cultured for an additional 3-7 days before being used for CAR expression analysis, in vitro coculture assay, and in vivo therapeutic experiments.

**In vitro coculture assay**

Vector control or SOAT1-KO firefly luciferase stably expressing Hepa1-6-OVA cells, as well as Vector control or SOAT1-KO HCC51-OVA cells were seeded in 96-well flat clear bottom white polystyrene TC-treated microplates (Corning, Cat#3610) at a concentration of $1\times10^4$ cells per well. After 2–4 h of tumor cell seeding, in vitro activated OT-I CD8+ T cells were added into tumor cells at Effector to Target ratios of 0.5:1, 1:1, or 2:1. After 24-72 h of coculture, the viability of Hepa1-6-OVA cells was assessed by measuring the remaining luciferase activity in tumor cells by adding 150 µg/ml D-Luciferin (PerkinElmer). After 48 h of HCC51-OVA and OT-I T cell coculture, the cells were stained with Fixable Viability Dye eFluor™ 506 (Thermo, Cat# 65-0866-14) and anti-mouse CD45. Tumor cell viability was assessed by quantifying percentages of viable CD45- cells with flow cytometry.

For in vitro CAR-T killing experiments, Vector control or SOAT1-KO luciferase stably expressing Huh7 cells, as well as Vector control or SOAT1-KO HepG2 cells were seeded in 96-well flat clear bottom white polystyrene TC-treated microplates (Corning, Cat#3610) at a concentration of $1\times10^4$ cells per well. After 2–4 h of cancer cell seeding, GPC3-targeting CAR-T cells were added into these cancer cells at Effector to Target ratios of 0.5:1, 1:1, or 2:1. The viability of HepG2 cells were evaluated by flow cytometry as used for HCC51-OVA cells.

To assess the tumor cell and T cell status by flow cytometry, cells were collected after 48 h of coculture and washed twice with MACS buffer. Cell surface marker such as anti-CD45, anti-CD8α was stained for 30 min at 4°C before lipid ROS staining, fillipin staining, and other intracellular cytokine staining.

**Liver tumor growth in C57BL/6 and Rag2$^{-/-}$ mice.**

To study the influence of SOAT1 knockout on liver tumor growth, SOAT1-KO or vector-transduced Hepa1-6 cells were subcutaneously transplanted into 6-8-week-old female C57BL/6 mice at a dose of $5\times10^6$ cells per mouse or $Rag2^{-/-}$ mice at a dose of $2\times10^6$ cells per mouse. Tumor sizes were monitored every 3–4 days using calipers by measuring tumor length and width.

For in vivo T cell therapy study, SOAT1-KO Hepa1-6-OVA cells or vector control Hepa1-6-OVA cells were subcutaneously transplanted into 6-8-week-old female $Rag2^{-/-}$ mice at a dose of $2\times10^6$ cells per mouse. 14 days after tumor inoculation, $0.5\times10^6$ of in vitro activated OT-I CD8+ T cells were intravenously transferred into tumor-bearing $Rag2^{-/-}$ mice. Therapeutic efficacy was assessed by monitoring tumor growth.

For anti-PD1 treatment study, syngeneic liver tumors were established by subcutaneously transplanting SOAT1-KO or vector control Hepa1-6 cells into 6-8-week-old C57BL/6 mice at a dose of $5\times10^6$ cells per mouse. Three doses of 8 mg/kg anti-PD1 were intraperitoneally administered into tumor-bearing C57BL/6J mice every 3-day starting from day 10 or day 14 when the average tumor volume reaches 300-500mm$^3$. For the combinatorial use of SOAT1 inhibitor, avasimibe, and anti-PD1, syngeneic liver cancer were established by subcutaneous transplanting $5\times10^6$ of Hepa1-6 cells into C57BL/6 mice. 15 mg/kg avasimibe (daily) and 8 mg/kg anti-PD1 (every 3-day) were subcutaneously administrated into tumor-bearing mice starting from 14 days post-tumor inoculation. Therapeutic efficacy was assessed by monitoring tumor growth.

For the study of tumor growth under the treatment of avasimibe, T cell therapy or their combination, SOAT1-KO or vector control Huh7 cells were subcutaneously transplanted into the flanks of 6-8 week-old female B-NDG mice at a dosage of $2\times10^6$ cells per mouse. One week post-tumor inoculation, avasimibe was administered via intraperitoneal injection at a dosage of 15 mg/kg daily. Ten days after tumor inoculation, PBS, avasimibe, $2\times10^6$ CART cells, or their combination were intravenously transferred into tumor-bearing B-NDG mice. The therapeutic efficacy was assessed by monitoring tumor growth.

Tumor sizes were calculated using the formula: Vol = 1/2×length×width^2. The statistical significance of all tumor growth curves in the present study was assessed using two-way analysis of variance (ANOVA), jointly considering the effect of treatment and the passage of time on tumor growth.

**Liver cancer growth under immunosuppressive conditions**

To study the effects of different immunosuppressants on liver cancer growth, we transplanted $5\times10^4$ Hepa1-6 cells subcutaneously into C57BL/6 mice treated with tacrolimus (1 mg/kg/day), sirolimus (1 mg/kg/day), sequential treatment with tacrolimus (7 days) followed by sirolimus, or PBS starting on the day of tumor inoculation. To evaluate the impact of SOAT1 inhibition on cancer recurrence and growth under normal and immunocompromised conditions, we transplanted $0.2\times10^6$ or $1\times10^6$ SOAT1-KO or vector control Hepa1-6 tumor cells subcutaneously into mice treated with PBS, or tacrolimus (1 mg/kg/day) 3 days before tumor inoculation. For the experiments using avasimibe to inhibit SOAT1, avasimibe was administered together with tacrolimus at a dose of 15 mg/kg/day.

**Orthotopic liver cancer study**

To evaluate orthotopic liver cancer growth and recurrence, $0.5\times10^5$ of SOAT1-KO and vector-transduced Hepa1-6 tumor cells were either injected subcapsularly into the liver or intrasplenically into immunocompetent C57BL/6 mice or immunodeficient Rag2$^{-/-}$ mice. Tumor progression was periodically monitiered by measuring luciferase activity using an intravital imaging system (IVIS). The procedure was as follows: 30 mg/mL firefly D-Luciferin potassium salt in saline was intraperitoneally administered to the experimental mice at a dose of 150 mg/kg body weight (typically 100-150 μL). The mice were anesthetized with isoflurane, and imaged using the IVIS system (PerkinElmer) after 10-15 min of D-Luciferin injection. The acquired images were analyzed using Living Image software to assess liver cancer size and progression. Mice were monitored using the Body Condition Scoring (BCS) system. Euthanasia was performed when their body condition severely deteriorated (BCS < 2). The date of death or euthanasia was recorded as the final survival day. Survival curves were generated using the Kaplan-Meier method and statistical significance was assessed by log-rank test.

**RNA extraction, quantitative RT-PCR, and RNA-seq**

RNA extraction from the target cells was performed using TRIzol Reagent (Invitrogen) according to the manufacturer's standard protocol. For RNA-seq analysis, the purified mRNA was utilized to prepare sequencing libraries with the NEBNext Ultra II Directional RNA Library Prep Kit for Illumina (NEB). These libraries were then multiplexed and sequenced using paired-end reads on Illumina Novaseq platform. Differential gene expression, Gene Set Enrichment Analysis (GSEA), and data visualization were conducted using R (v.5.2.0), and the ggplot2 (v3.2.1). For quantitative RT-PCR (qPCR) analysis, first-strand cDNA was synthesized from extracted RNA using the HiScript Q RT SuperMix for qPCR (Vazyme) or the HiScript III 1st Strand cDNA Synthesis Kit (Vazyme). The resulting cDNA was diluted and normalized with nuclease-free water before being used for qPCR. Quantitative PCR was then conducted with Taq Pro Universal SYBR qPCR Master Mix and gene-specific primers, using GAPDH or ACTB as an internal control.

**Flow cytometry analysis and FACS sorting.**

Cells from different treatment groups were collected, washed twice, and resuspended in 50-100 μL fluorophore-conjugated antibodies-containing MACS buffer at a density of 0.1-1×$10^7$ cells per ml. The cells were stained at 4°C for 30 min. Following staining, cells were washed twice with MACS buffer before being used for flow cytometry analysis or cell sorting. For intracellular staining, cells were treated with 5 μg/ml Brefeldin A and PMA/Ionomycin, incubated for 4 h, then collected, washed and stained with antibodies for cell surface markers. After surface staining, the cells were washed twice and fixed in 4% paraformaldehyde at 4°C for 15-20 min. The fixed cells were permeabilized using 0.1% Triton-X100 or eBioscience permeabilization buffer for 10-15 min at room temperature, then washed twice. Subsequently, the cells were stained with antibodies against intracellular proteins for 30 min on ice. Finally, the cells were washed, resuspended in MACS buffer, and subjected to flow cytometry analysis.

**Flow analysis of spleen and tumor immune microenvironment**

At the study endpoint, tumor-bearing mice were euthanized, and tumors along with draining lymph nodes (dLN) were harvested and placed in ice-cold FACS buffer (2%FBS in PBS). Lymph nodes were directly processed by smashing and filtering through 100 μm cell strainer to obtain single cell suspension. Tumors were minced into small pieces using a scraper, and digested in DMEM containing 100 U/mL Collagenase IV at 37°C for 30-60 min. Following digestion, tumor suspensions were diluted with PBS and filtered through 100 μm cell strainer to generate single-cell suspension. The single cell suspension from tumors was washed twice and then treated with 3-5 ml of RBC Lysis Buffer at room temperature for 5-10 min to lyse red blood cells. After lyses, the cells was diluted with 10 volumes of FACS buffer and centrifuged at 400 g for 5 min at 4°C to pellet the cells. Cells from both tumors and dLN were resuspended and counted for flow cytometry staining as detailed in the "Flow cytometry analysis and FACS sorting" section. Data were analyzed using FlowJo software (v.9.9.4 or v.10.3). Immune cell subpopulations, including CD4+T cells, CD8+T cells, monocytes, neutrophils, dendritic cells and macrophages, were identified using the previously reported gating strategy[3].

**Cellular Lipid ROS, total antioxidant capacity, GSH and GSSG detection**

To detect lipid ROS, cells from different treatment group in vitro and in vivo, were stained with 10 μM BODIPY581/591C11 dye solution at 37°C for 30 min. Then the cells were washed twice with MACS buffer and then incubated with flow antibody at 4°C for 30 min. After washing twice with MACS buffer, cells were analyzed by flow cytometry. The mean fluorescence intensity of FITC and PE-Texas Red within the desired cell populations was calculated, and the levels of oxidized lipids was assessed by calculating the FITC/ PE-Texas Red ratio. Total cellular antioxidant capacity and oxidative status were evaluated using

Total Antioxidant Capacity Assay Kit with ABTS method (Beyotime, S0121) and GSH/GSSG ratio detection kit (Beyotime, S0053), by following manufacturer's instructions.

**Single cell RNA-seq analysis**

Fresh tumor tissue samples were cut into approximately 1mm$^3$ pieces and washed three times with 4 ml of cold PBS. The samples were then minced into small pieces, and enzymatically digested using the MACS Tumor Dissociation Kit (Miltenyi Biotec) with agitation for 30 min, according to the manufacturer's instructions. The dissociated cell suspension were filtered sequentially through a 70-µm and 40-µm cell-strainers, and centrifuged at 300 g for 10 min. After centrifugation, the supernatant was discarded, and the cell pellet was resuspended in red blood cell lysis buffer (Tiangen Biotech#RT122-1) and incubated at room temperature for 5 min to lyse red blood cells. The cells were then washed twice with washing buffer (0.04% bovine serum albumin in PBS) and resuspended in the same buffer. The cells were counted using Countess™ 3 Automated Cell Counter (Life), and the final concentration was adjusted to 1,000 cells/µL usually with a viability of ≥80%.

For single-cell RNA sequencing (scRNA-seq) library preparation, the DNBelab C Series High-throughput Single-Cell RNA Library Prep Kit (MGI, Cat# 940-000519-00) was used. In brief, cells at a concentration of 1,000 cells/µL were loaded into the cell reservoir of a microfluidic chip. Barcoded beads and droplet-generation oil were sequentially added to their respective reservoirs. Encapsulated droplets were generated and collected using the DNBelab C4 /DNBelab TaiM4 system. Beads capturing the mRNA were then recovered for reverse transcription (RT). After RT, complementary DNA (cDNA) was amplified via PCR, purified, and quantified using a Qubit dsDNA kit (Thermo Fisher). Libraries of 3'-end transcripts were subsequently constructed through cDNA fragmentation, size selection, end repair and A-tailing, adapter ligation, indexing PCR, and library cyclization, according to the manufacturer 's protocol. The sequencing libraries were purified and quantified using the Qubit ssDNA kit (Thermo Fisher) and the Qsep100 system (Bioptic). scRNA-seq was performed on the DNBelab C4/DNBelab TaiM4 system. The DNBelab C4 /DNBelab TaiM4 Series Single-Cell Library Prep Set (MGI) was used for sequencing. DNBs were loaded into patterned nanoarrays and sequenced on the DNBSEQ-T7 sequencer with pair-end sequencing. The sequencing reads contained a 30-bp read 1 (including a 10-bp cell barcode 1, a 10-bp cell barcode 2 and a 10-bp unique molecular identifiers (UMI)), a 100-bp read 2 for gene sequences, and a 10-bp barcode read for sample indexing.

Sequencing data were processed using the open-source DNBelab C Series scRNA analysis software pipeline (https://github.com/MGI-tech-bioinformatics/DNBelab_C_Series_scRNA-analysis-software).

Sample de-multiplexing, barcode processing, and single-cell 3' UMI counting were performed with default parameters. Processed reads were aligned to the GRCh38 genome reference using STAR (2.7.2b). Valid cells were automatically identified based on UMI distribution using the "barcodeRanks" function of the DropletUtils tool to remove background beads and low-UMI-count beads. Finally, we used PISA tool to calculate gene expression for each cell and create a gene-by-cell matrix for each library.

**Multiplex immunofluorescence assay**

After dewaxing of the paraffin sections in water, the slides were immersed in a citric acid antigen retrieval buffer and boiled for antigen retrieval. After cooling, the samples were incubated with 3% hydrogen peroxide ($H_2O_2$) at room temperature to quench endogenous peroxidase activity, followed by rinsing with PBS. The sections were then blocked with 5% FBS at 37 °C for 30 min to prevent nonspecific binding. After blocking, the sections were incubated with the primary antibody overnight at 4°C. The next day, the slides were rinsed with PBS and incubated with the appropriate secondary antibodies at 37 °C, followed by another PBS rinse. Subsequently, the sections were stained with Hoechst at room temperature for 15 min to visualize nuclei. For serial staining, the primary and secondary antibodies were stripped off using microwave treatment, and the process was repeated as needed for additional staining.

**Data statistics.**

Standard non-NGS data analysis was conducted using GraphPad Prism10. NGS statistical analyses were performed in R/RStudio. Unless specified, a two-sample unpaired t-test was used to compare two groups. Tumor growth curves were analyzed by a two-way ANOVA with two independent variables. In clinical samples, propensity score matching analysis was conducted among 16 obese patients based on tumor stage, pathological grade, and age, resulting in an equal number of baseline-matched normal patients.
